# Phospholipase A_2_ inhibitor-loaded micellar nanoparticles attenuate inflammation and mitigate osteoarthritis progression

**DOI:** 10.1101/2021.01.15.426857

**Authors:** Yulong Wei, Lesan Yan, Lijun Luo, Tao Gui, Ahmad Amirshaghaghi, Tianyan You, Andrew Tsourkas, Ling Qin, Zhiliang Cheng

**Affiliations:** Department of Bioengineering, School of Engineering and Applied Sciences, University of Pennsylvania, Philadelphia, Pennsylvania 19104, USA; Department of Orthopaedic Surgery, Perelman School of Medicine, University of Pennsylvania, Philadelphia, Pennsylvania 19104, USA; Department of Orthopaedics, Union Hospital, Tongji Medical College, Huazhong University of Science and Technology, Wuhan 430022, China; School of Agricultural Equipment Engineering, Jiangsu University, Zhenjiang, Jiangsu 212013, China

## Abstract

Treating osteoarthritis (OA) remains a major clinical challenge. Despite recent advances in drug discovery and development, no disease-modifying drug for knee OA has emerged with any significant clinical success, in part due to the lack of valid and responsive therapeutic targets and poor drug delivery within knee joints. In this work, we show that the amount of secretory phospholipase A_2_ (sPLA_2_) enzyme increases in articular cartilage in human and mouse OA cartilage tissues. We hypothesize that inhibition of sPLA_2_ activity may be an effective treatment strategy for OA. To develop a sPLA_2_-responsive and nanoparticle (NP)-based interventional platform for OA management, we incorporated a sPLA_2_ inhibitor (sPLA_2_i) into the phospholipid membrane of micelles. The engineered sPLA_2_i-loaded micellar nanoparticles (sPLA_2_i-NPs) were able to penetrate deep into the cartilage matrix, prolong retention in the joint space, and mitigate OA progression. These findings suggest that sPLA_2_i-NPs can be promising therapeutic agents for OA treatment.

## INTRODUCTION

Osteoarthritis (OA) is a painful and debilitating disease of articular cartilage, leading to joint pain, loss of joint function, and deleterious effects on the quality of daily life. It occurs in approximately 27 million adults in the United States alone, with staggering societal and economic costs (60 billion/year) (*1*). Current treatments for OA include non-pharmacological treatment (e.g., diet or exercise), pharmacological approaches and surgical intervention (*2*). Although many pharmacologic approaches have been extensively pursued and some drugs have shown promise in preclinical studies, none has emerged with any significant clinical success, and there are no disease-modifying therapies available to delay and/or limit OA development and progression (*3*). Accordingly, there is a great unmet medical need to develop new interventional platforms for greater effectiveness in OA treatment.

The etiology of OA is broad and includes various mechanical, biochemical, and genetic factors (*4, 5*). Recent work has indicated that chronic unresolved inflammation has a critical role in OA development and progression (*6, 7*). Analyses of tissues from both human patients and animal models of OA have revealed that different inflammatory mediators have been implicated in OA pathogenesis (*4*). However, the most widely used nonsteroidal anti-inflammatory agents (NSAIDs) only provide the short-term management of the pain symptoms in OA. Moreover, they have substantial renal and cardiovascular toxicities and limited efficacy without repeated administration (*8*). Thus, the identification of new anti-inflammatory targets may provide benefits in the treatment of OA disease.

Among the potent inflammatory mediators involved in the development of OA is the family of secreted phospholipase A_2_ (sPLA_2_) enzymes. sPLA_2_ is a heterogeneous group of enzymes that specifically recognize and catalytically hydrolyze the sn-2 ester bond of glycerophospholipids, releasing free fatty acids such as arachidonic acid (AA) and lysophospholipids (*9*), that are well-known mediators of inflammation and tissue damage (*10–16*). sPLA_2_ is normally present at low levels in healthy knee joint tissues. Yet, under pathological conditions sPLA_2_ can be induced by multiple cascades and effector molecules including inflammatory cytokines (*17, 18*) and free radicals (*19*). A high expression level and activity of sPLA_2_ is observed in the synovial membrane, synovial fluid, and articular cartilage of human OA patients (*20–23*). As such, we postulated that inhibition of sPLA_2_ enzyme activity by sPLA_2_i can be exploited as a novel therapeutic strategy for OA treatment. Note that, several sPLA_2_i compounds have already been developed and used in clinical trials for treating other inflammatory diseases (*24*). However, few studies have examined the role of sPLA_2_ in OA and no study has sought to harness the sPLA_2_ activity for the treatment of OA (*18*).

Clinical management of OA is further complicated by lack of effective drug delivery systems (*25, 26*). Rapid clearance of the drugs by the joint (i.e. a short half-life) and therapeutic target sites deep within the cartilage not accessible to drugs pose significant delivery challenges for many promising drugs (*27, 28*). Recently, nanoparticle-based targeted drug delivery has been exploited in the treatment of OA (*1, 27, 29*). The main benefits of these delivery systems are that they can increase the retention of drugs in the joints, thus lowering the therapeutic dose, reducing the frequency of administration, increasing therapeutic efficacy, and reducing off-target toxicity. Here, we examined whether inhibition of sPLA_2_ enzyme activity by sPLA_2_i-loaded nanoparticles could be an effective OA therapy. Specifically, a lipid-based sPLA_2_i, thioetheramide-PC, was incorporated into nanometer-sized phospholipid micelles. The joint retention, cartilage penetration, biodistribution and toxicity of the sPLA_2_i-loaded micellar nanoparticles (sPLA_2_i-NPs) were subsequently characterized. Using an in vitro cartilage explant model and two animal models of OA, we investigated the effectiveness of sPLA_2_i-NPs in inhibiting the inflammatory signals and attenuating the OA progression following direct delivery into knee joints.

## RESULTS

### sPLA_2_ amount in OA cartilage

To confirm the elevated levels of sPLA_2_ enzyme in OA-related tissues, we performed immunohistochemistry (IHC) to investigate a commonly studied isoform sPLA_2_-IIA in OA and normal knees. In healthy young and adult human cartilage, the level of sPLA_2_-IIA was very low (Fig. 1a,b). However, in OA cartilage, it was drastically elevated after OA initiation and remained at a high level at early-, middle-, and late-stages of OA progression. Abundant staining was found in the cartilage matrix as well as chondrocytes. Mouse cartilage showed similar results (Fig. 1c,d). While normal mouse articular cartilage displayed only a weak staining of sPLA_2_-IIA, cartilage in mouse knees receiving destabilization of the medial meniscus (DMM) surgery a month earlier showed greatly elevated staining throughout the cartilage. The marked increase of sPLA_2_ in human and mouse OA cartilage suggested its potential role in OA development.

**Fig. 1.**
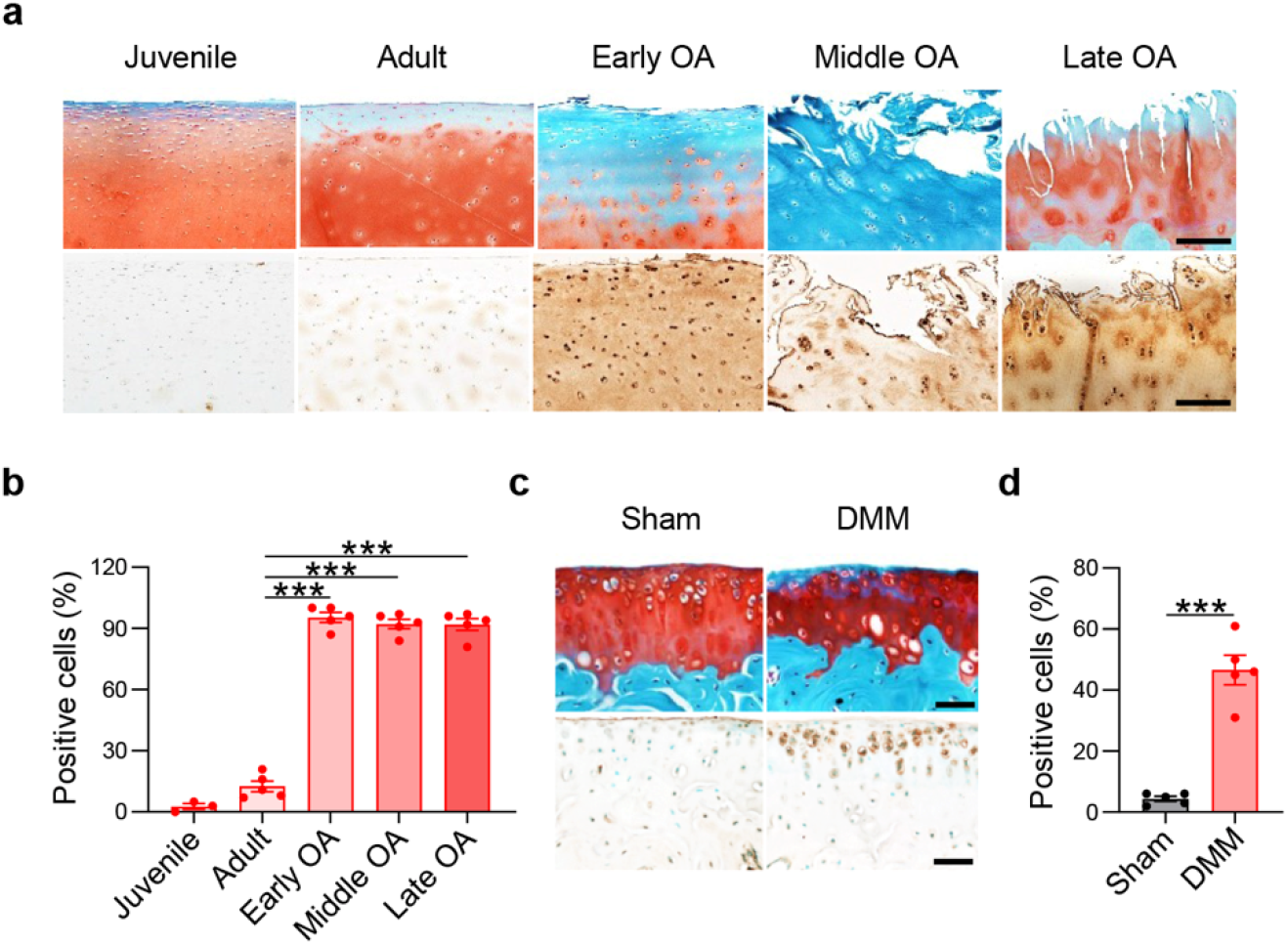
sPLA_2_ expression in human and mouse OA cartilage. **a,** Representative images of safranin O/fast green staining (top) and immunohistochemistry of sPLA_2_ (bottom) in healthy juvenile, healthy adult, early stage, middle stage and late stage OA human cartilage tissues. Scale bars, 200 μm. **b,** Quantification of sPLA_2_-positive chondrocytes as a proportion of total chondrocytes in healthy and OA human articular cartilage tissues. (n = 5). **c,** Representative images of safranin O/fast green staining (top) and immunohistochemistry of sPLA_2_ (bottom) in tibial articular cartilage of *WT* mice at 1 month post-sham or post-DMM surgery. Scale bars, 50 μm. **d,** Quantification of sPLA_2_-positive chondrocytes in tibial articular cartilage of sham- or DMM-operated joints (n = 5). Statistical analysis was performed using one-way ANOVA with Dunnett’s post hoc test for (b) and paired two-tailed t-test for (d). Data presented as mean ± s.e.m. ***p<0.001.

### sPLA_2_i-NP synthesis and characterization

sPLA_2_i-NPs were prepared by incorporating 25 mol% thioetheramide-PC into phospholipid micelles with lipid composition 10□mol% 1,2□dioleoyl□3□trimethylammonium propane (DOTAP)/65□mol% 1,2□distearoyl□sn□glycero□3□phosphoethanolamine□N□[methoxy(polyethylene glycol)□2000] (mPEG2000□DSPE) (Fig. 2a). Due to its amphiphilic nature, lipid-based sPLA_2_i was easily doped into the phospholipid film during nanoparticle preparation. Dynamic light scattering (DLS) measurements revealed that sPLA_2_i-NPs possessed a mean hydrodynamic diameter of 10 nm and a relatively narrow size distribution (Fig. 2b). sPLA_2_i-NPs observed by transmission electron microscopy (TEM) were approximately spherical in shape together with some worm-like structures. As shown in Fig. 2c, the presence of cationic DOTAP within sPLA_2_i-loaded nanoparticles transitioned nanoparticles from a negative surface charge (−8 mv) to a positive surface charge (+2 mv), to increase the retention and penetration ability of sPLA_2_i-NPs within the cartilage via electrostatic interactions between the nanoparticles and the anionic glycosaminoglycans (GAGs) in the cartilage. The stability of the sPLA_2_i-NPs was evaluated in water and bovine synovial fluid. There was no observable change in the hydrodynamic diameter of sPLA_2_i-NPs in water for at least 1 week (Fig. 2d) or in bovine synovial fluid for 24 hours (Supplementary Fig. S1).

**Fig. 2.**
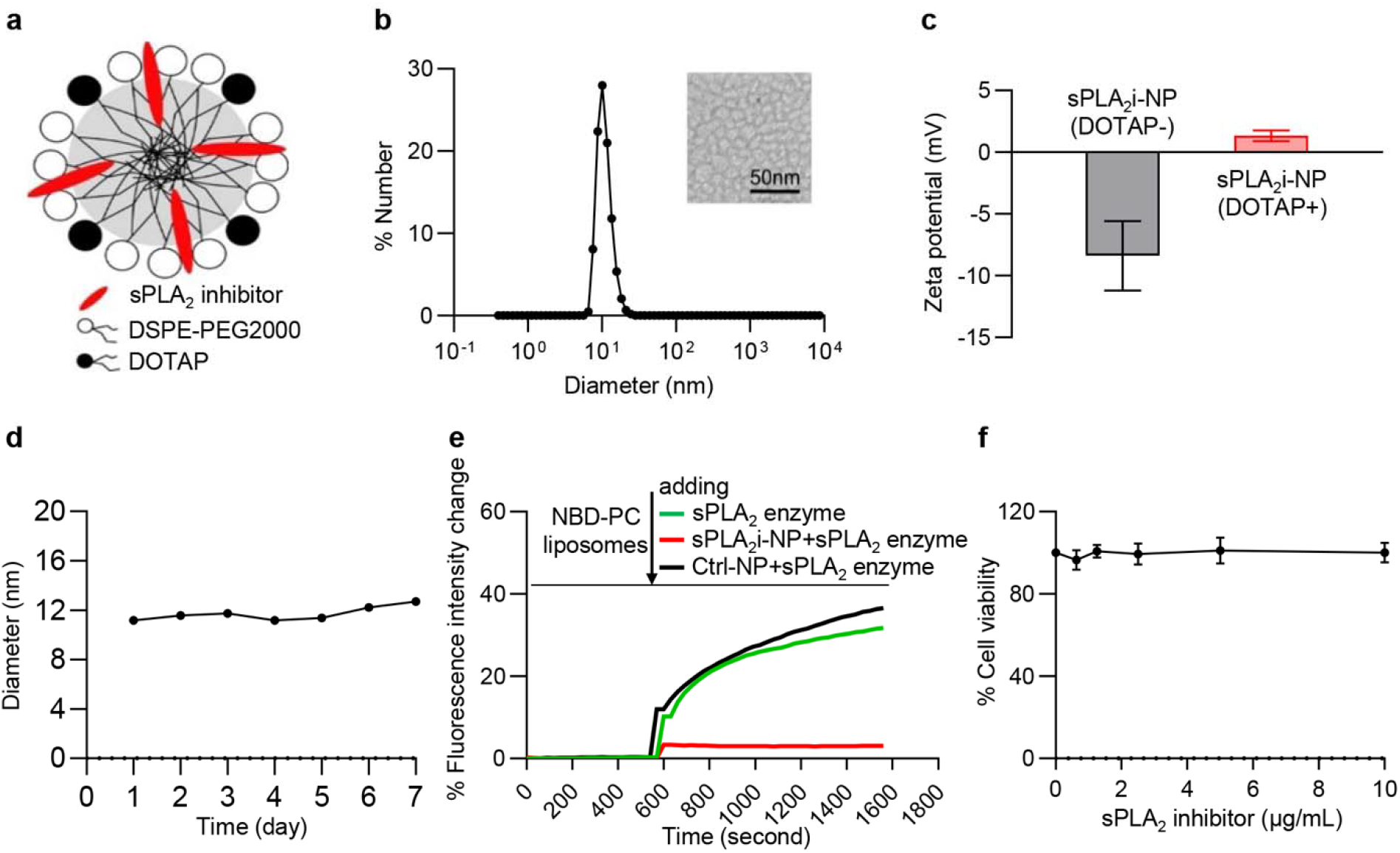
Preparation and characterization of sPLA_2_i-NPs. **a,** Schematic diagram of sPLA_2_i-NPs, in which a lipid-based sPLA_2_i was incorporated into nanometer-sized phospholipid nanoparticles (DSPE-PEG2000) with cationic lipid DOTAP doped. **b,** DLS measurement of sPLA_2_i-NPs hydrodynamic diameter and TEM (insert) of sPLA_2_i-NPs. **c,** Zeta potential of sPLA_2_i-NPs in the presence or absence of cationic lipid DOTAP in 0.1 M PBS (pH = 7.4). **d,** The stability of sPLA_2_i-NPs in water was accessed by monitoring the hydrodynamic diameter for up to 1 week. **e,** In vitro response of sPLA_2_i-NPs to sPLA_2_ enzyme experiment showed a significant inhibition effect. **f,** The cytotoxicity of sPLA_2_i-NPs was determined by measuring the cell viability of primary chondrocytes after co-incubation with sPLA_2_i-NPs at various concentrations. In all datasets, n=3 biologically independent experiments. Statistical analysis was performed using one-way ANOVA with Dunnett’s post hoc test. Data presented as mean ± s.e.m.

To examine whether sPLA_2_i-NPs can inhibit sPLA_2_ enzyme, a fluorescence dequenching assay with liposomes that contain a self-quenching concentration of the fluorescent lipid was used(*30*). Phospholipid hydrogenated soy phosphatidylcholine (HSPC) liposomes doped with 20 mol% fluorescent lipid 1-palmitoyl-2-{6-[(7-nitro-2-1,3-benzoxadiazol-4-yl)amino]hexanoyl}-*sn*-glycero-3-phosphocholine (NBD-PC) were prepared in which the NBD fluorescence was self-quenched (Supplementary Fig. S2). No significant change in fluorescence intensity was observed when the preincubation mixture of sPLA_2_i-NPs and sPLA_2_ enzyme was added into NBD-incorporated liposomal suspension. However, a significant increase in the fluorescence intensity was observed upon the addition of the preincubation mixture of Ctrl-NPs (i.e., nanoparticles without sPLA_2_i) and sPLA_2_ enzyme (Fig. 2e). These results suggest that sPLA_2_i-NPs are sPLA_2_-responsive and can inhibit sPLA_2_ enzyme.

The cytotoxic effects of sPLA_2_i-NP were examined in a MTT cell proliferation assay. Specifically, various concentrations of sPLA_2_i-NPs were incubated with primary mouse chondrocytes for 24 hours. The cell viability for each group was normalized to a control group that was not incubated with any sPLA_2_i-NPs. In general, sPLA_2_i-NPs had little effect on the viability of cells up to a sPLA_2_i concentration of 10 ug/mL (Fig. 2f), suggesting their physiological biocompatibility.

### sPLA_2_i-NP penetration

To examine whether the sPLA_2_i-NPs could penetrate into deep layer of cartilage tissue, sPLA_2_i-NPs were incubated with bovine cartilage tissue ex vivo. Confocal fluorescence images of cartilage section were acquired preincubation and at various time points after incubation with sPLA_2_i-NPs (Fig. 3a). In the preincubation images, there was little intrinsic tissue fluorescence. At two days following incubation with the sPLA_2_i-NPs, a strong fluorescence signal was observed within the superficial zone of cartilage. The fluorescence signal was distributed in the entire cartilage tissue (1-2 mm thickness) as early as day 4, and became stronger over time, indicating the sPLA_2_i-NPs indeed moved into the deep zone of the cartilage. Control experiments to demonstrate the high penetration capability of sPLA_2_i-NP were performed by incubation of bovine cartilage explants with sPLA_2_i-NPs that were not doped with the positively-charged DOTAP. In these tissues, a strong fluorescence signal was only observed within the superficial zone of cartilage, even up to 8 days. Similarly, when bovine cartilage explants were incubated with free fluorescent dye rhodamine, very little fluorescence signal was observed in the deep zone of cartilage. Quantitative analysis of fluorescence images revealed that DOTAP-doped sPLA_2_i-NPs exhibited a great improvement in cartilage penetration at all time points compared with non-DOTAP doped sPLA_2_i-NPs or free rhodamine (Fig. 3b and Supplementary Fig. S3). In addition, the area under curve (AUC) in the cartilage achieved by sPLA_2_i-NPs (DOTAP+) was much larger than that of non-DOTAP doped sPLA_2_i-NPs or free rhodamine (Fig. 3c and Supplementary Fig. S3). These results indicate that the DOTAP-doped sPLA_2_i-NPs are able to penetrate into the deep zone of articular cartilage and exhibit high cartilage accumulation.

**Fig. 3.**
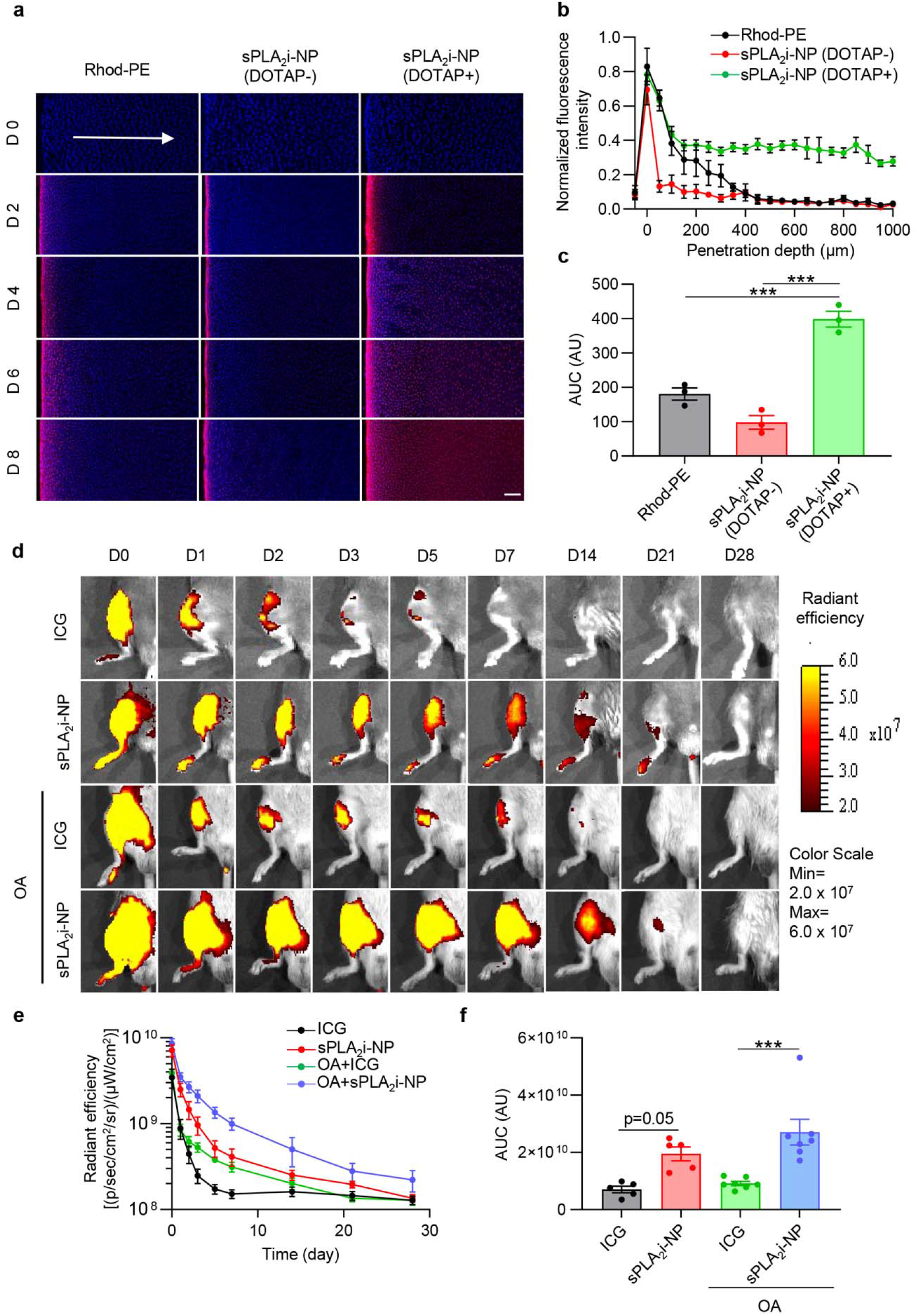
Cartilage penetration and joint retention of sPLA_2_i-NPs. **a,** Representative confocal microscope images of cross-sections of bovine cartilage explants incubated with free rhodamine dye or rhodamine-labeled sPLA_2_i-NPs in the presence or absence of cationic lipid DOTAP for 0, 2, 4, 6 and 8 days. Scale bar, 200 μm. **b,** Quantitative analysis of the fluorescence intensity of the above different formulations over the entire explant sections (n = 3). **c,** Quantification of the area under the curve (AUC) based on the fluorescence intensity profiles in b (n = 3). **d,** Representative IVIS images of healthy and OA (OA was induced surgically 8 weeks before) mouse knee joints over 28 days post single intra-articular injection of free ICG or Cy7-labeled sPLA_2_i-NPs. Fluorescent scale, min = 2.0 × 10^7^, max = 6.0 × 10^7^. **e,** Quantitative analysis of time course fluorescent radiant efficiency within healthy and OA mouse knee joints over 28 days (n = 5). **f,** Quantification of the area under the curve (AUC) based on the fluorescence intensity profiles in e (n = 5). Rhod-PE: free rhodamine dye, sPLA_2_i-NPs in the absence of DOTAP: sPLA_2_i-NPs (DOTAP-), sPLA_2_i-NPs in the presence of DOTAP: sPLA_2_i-NPs (DOTAP+). Statistical analysis was performed using one-way ANOVA with Turkey’s post hoc test. Data presented as mean ± s.e.m. ***p<0.001.

To further assess the total accumulation of sPLA_2_i-NPs within cartilage, fluorescence images of bovine cartilage were acquired 24 hours following the incubation of DOTAP-doped sPLA_2_i-NPs, non DOTAP-doped sPLA_2_i-NPs, or free rhodamine. As expected, the fluorescent intensity of DOTAP-doped sPLA_2_i-NPs in the cartilage tissue was significantly higher than that of non-DOTAP doped sPLA_2_i-NPs or free rhodamine (Supplementary Fig. S4a,b). This convincingly demonstrates the benefit of including DOTAP moieties in the sPLA_2_i-NPs. Due to the ability to penetrate cartilage, the DOTAP-doped sPLA_2_i-NPs were chosen for subsequent studies.

### sPLA_2_i-NP retention

The retention of Cy7-labeled sPLA_2_i-NPs in the knee joints post intra-articular injection was monitored using in vivo fluorescence imaging. To show the difference in retention between nanoparticles and small-molecule agents, a low molecular weight fluorescent dye indocyanine green (ICG, MW 775) was used for comparison. Their retention within the joints were evaluated in the knee under healthy and early OA conditions. Fluorescence images of mouse knee joints were acquired at various time points post sample injection (Fig. 3d). With a single intra-articular injection, the fluorescence intensity of sPLA_2_i-NPs in joints was significantly higher than that of free ICG at all time points, indicating that the retention time of sPLA_2_i-NPs in joints is much longer than that of free ICG. Quantitative analysis of fluorescence images showed that the retention of sPLA_2_i-NPs in joints with OA condition was even more efficient than that of healthy joints (Fig. 3e,f). A longer retention time of sPLA_2_i-NPs over small molecules within knee joints was also confirmed when rats were used in this study (Supplementary Fig. S5).

We also examined the biodistribution of sPLA_2_i-NPs in joint components, internal organs, and blood. At 24 hours post injection, fluorescence signals were detected on the cartilage surfaces of patellar, femur condyles and tibiae plateau as well as on the meniscus (Supplementary Fig. S6a,b). By 24h postinjection, the accumulation of the sPLA_2_i-NPs was mainly observed in the liver and kidney, but no signal was detected in blood at that time, indicating sPLA_2_i-NPs were nearly cleared from circulation (Supplementary Fig. S7a,b). One month later, no fluorescence was observed in the liver and kidney.

### sPLA_2_i-NPs block cartilage degeneration in OA cartilage explants

Articular cartilage explants represent an in vitro model to study OA progression in a three dimensional (3D) environment(*31*). We harvested femoral head cartilage explants from 2-month-old wild-type (*WT*) mice and then stimulated them with OA-associated pro-inflammatory cytokine interleukin-1β (IL-1 β)(*32*). The penetration of sPLA_2_i-NPs was studied by acquiring fluorescence images of IL-1β-stimulated cartilage explants that were incubated with rhodamine-labeled sPLA_2_i-NPs for 24 or 48 hours. Fluorescent signal was mainly present in the cartilage surface at 24 hours (Fig. 4a), while a strong fluorescence signal was observed in the deep calcified layer at 48 hours (Fig. 4b,c), indicating that sPLA_2_i-NPs are able to penetrate into femoral head cartilage treated with IL-1β. These results were consistent with those presented above for sPLA_2_i-NPs penetration within bovine cartilage explants.

**Fig. 4.**
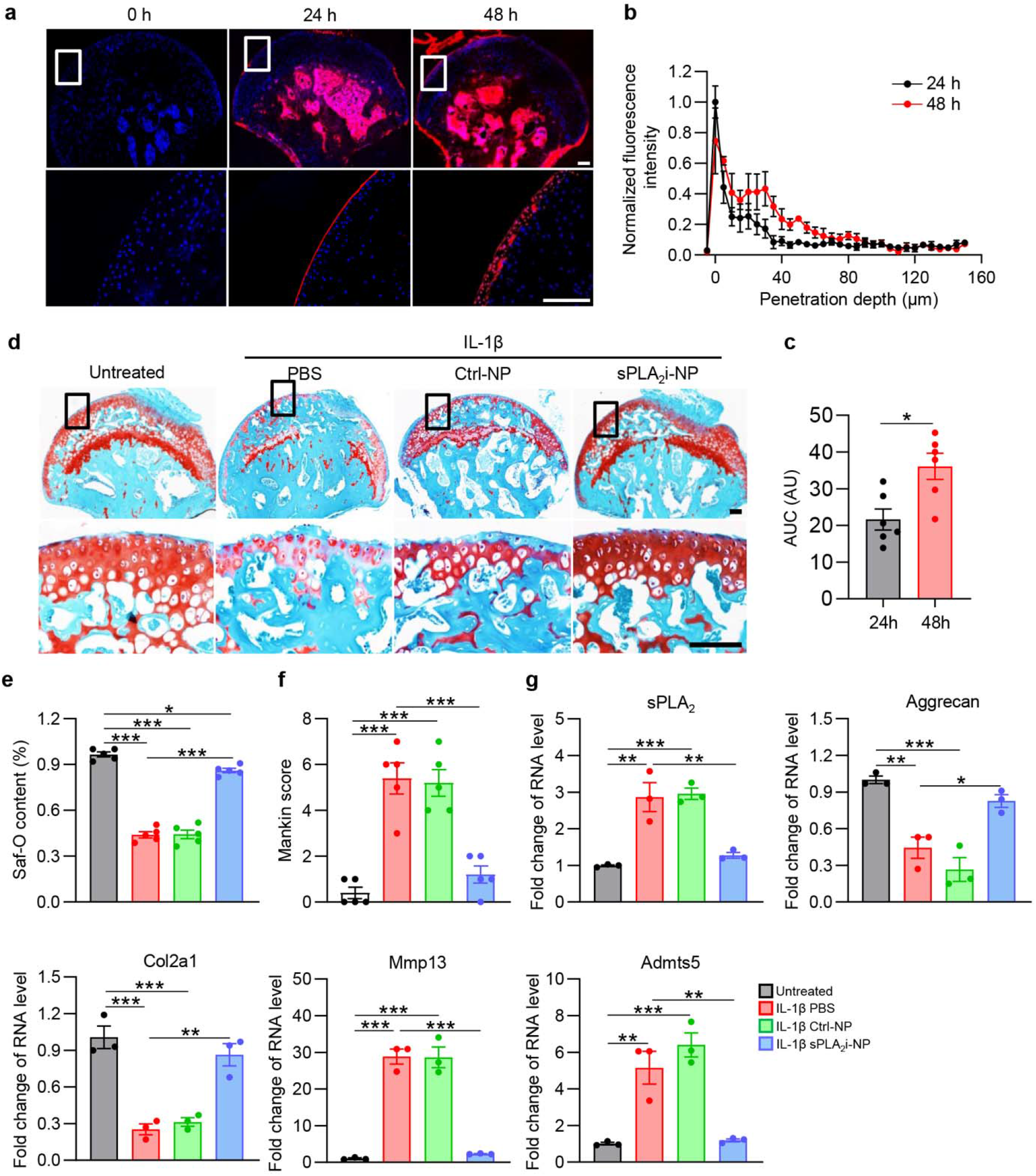
Penetration and chondroprotective effects of sPLA_2_i-NPs in mouse femoral heads. **a,** Representative fluorescence images of cross-sections of mouse femoral heads stimulated with recombinant mouse IL-1β and incubated with rhodamine-labeled sPLA_2_i-NPs for 0, 24 or 48 hours. Magnified images of the white boxed areas are presented as bottom panels. Scale bars, 100 μm. **b,** Quantitative analysis of rhodamine-labeled sPLA_2_i-NPs penetration depth into IL-1β-stimulated mouse femoral heads (n = 6). **c,** Quantification of the area under the curve (AUC) based on the fluorescence intensity profiles in b (n = 6). **d,** Representative images of safranin-O/fast green staining on the sections of untreated and PBS-, Ctrl-NP- and sPLA_2_i-NP-treated IL-1β-stimulated mouse femoral heads. Magnified images of the black boxed areas are presented as bottom panels. Scale bars, 100 μm. **e,** Safranin-O-positive area among the above groups was quantified (n = 5). **f,** The OA severity was accessed by Mankin score (n = 5). **g,** The relative gene expression of sPLA_2_-IIA, Aggrecan, Col2a1, Mmp13 and Adamts5 was examined by qRT-PCR in the untreated and PBS-, Ctrl-NP- and sPLA_2_i-NP-treated IL-1β-stimulated mouse femoral heads (n = 3). Statistical analysis was performed using paired two-tailed t-test for (c) and one-way ANOVA with Turkey’s post hoc test for (e), (f) and (g). Data presented as mean ± s.e.m. *p<0.05, **p<0.01, ***p<0.001.

To test the therapeutic effects of sPLA_2_i-NPs in vitro, femoral head cartilage explants were divided into 4 groups that received PBS, IL-1β and PBS, IL-1β and Ctrl-NPs, or IL-1β and sPLA_2_i-NP treatments, respectively, for 8 days. As shown in Fig. 4d-f, IL-1β treatment led to an OA-like phenotype in cartilage, featured by surface fibrillation and proteoglycan loss. Strikingly, sPLA_2_i-NPs, but not Ctrl-NPs, blocked cartilage degeneration induced by IL-1β, leading to comparable Safranin-O content and Mankin score as the untreated group. sPLA_2_i-NPs, but not PBS or Ctrl-NPs, attenuated IL-1β-induced sPLA_2_-IIA expression in cartilage (Fig. 4g). Furthermore, the catabolic effects of IL-1β, such as decreasing the expression of matrix proteins, Col2a1 and Aggrecan, and increasing the expression of proteinases, Mmp13 and Adamts5, were effectively reversed by sPLA_2_i-NPs. (Fig. 4g). Taken together, these data provide ex vivo evidence that sPLA_2_i-NPs have protective action on chondrocytes against OA inducing insults.

### sPLA_2_i-NPs attenuate joint destruction in a surgery-induced mouse OA model

To study the in vivo therapeutic effects of sPLA_2_i-NPs, we used two mouse OA models. The first, a DMM surgery model, mimics chronic OA development in human patients (Fig. 5a). After DMM, *WT* knees started to show cartilage damage, including surface fibrillation and loss of proteoglycan, at 2 months, and exhibited severe cartilage erosion up to the entire uncalcified zone at 4 months, resulting in Mankin scores of 5.5 and 8.9, at these two time points respectively (Fig. 5b,c and Supplementary Figs. S8 and S9). Intra-articular injection of sPLA_2_i-NPs into DMM knees, once every two weeks, starting immediately after the surgery, greatly improved the morphology and structure of articular cartilage, leading to an almost intact cartilage surface with no proteoglycan loss even at 4 months post-surgery (Fig. 5b and Supplementary Figs. S8 and S9b). These therapeutic effects were not observed with free sPLA_2_i or Ctrl-NP treatments.

**Fig. 5.**
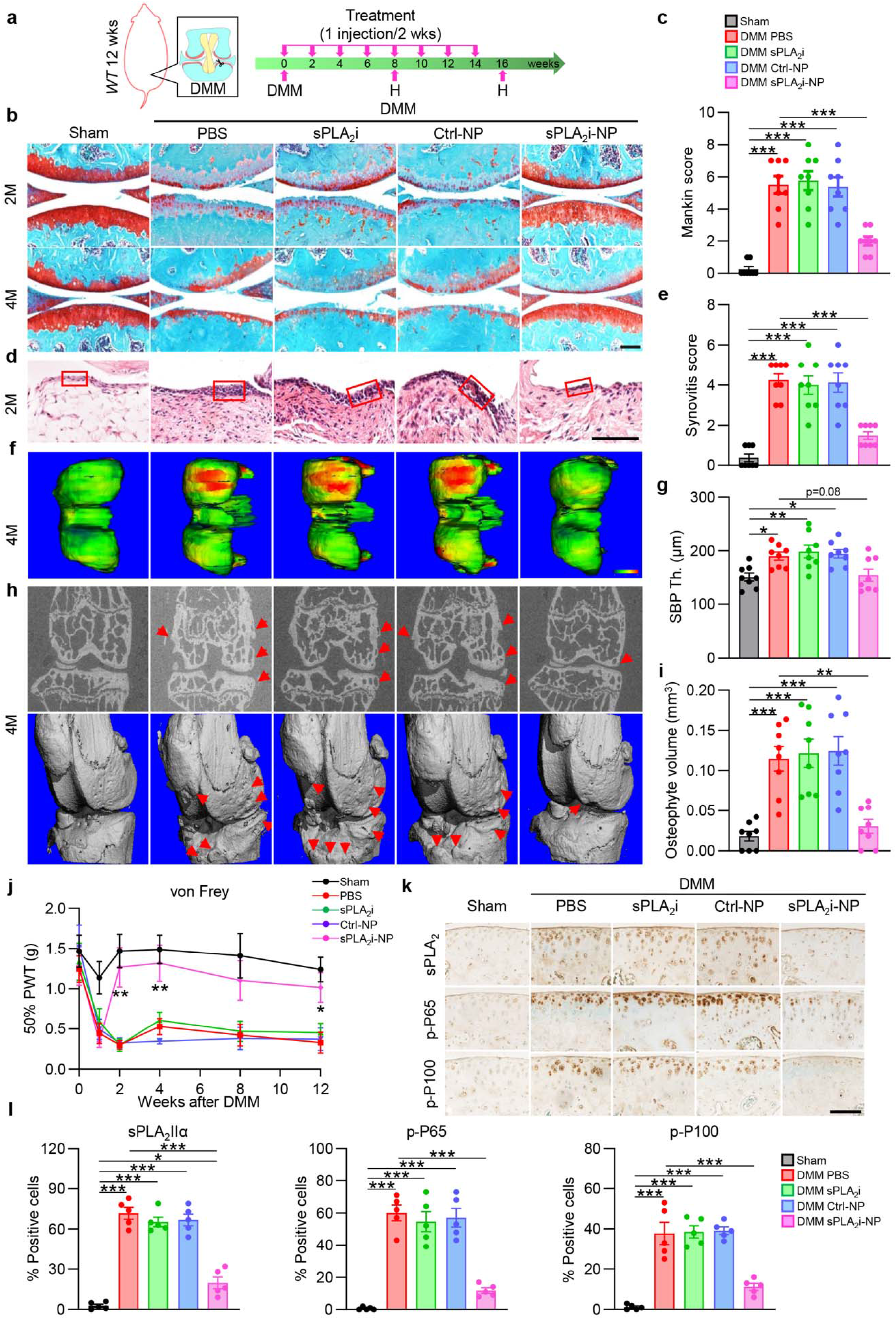
Therapeutic effects of sPLA_2_i-NPs for attenuation of DMM-induced traumatic OA. **a,** The study design of sPLA_2_i-NP treatment for DMM-induced OA mice. DMM surgery was performed on 3-month-old male mice followed by intra-articular injections of PBS, sPLA_2_i, Ctrl-NPs and sPLA_2_i-NPs once every week for 2 or 4 months. **b,** Representative images of safranin O/fast green staining in the articular cartilage of sham- or DMM-operated knee joints with 2- or 4-month treatment. Scale bar, 200 μm. **c,** The OA severity was accessed by Mankin score after 2-month treatment (n = 8). **d,** Representative images of H&E staining in the sham- or DMM-operated knee joints with 2-month treatment. Scale bar, 100 μm. Red boxed area indicate the enlargement of synovial lining cell layer. **e,** Synovial inflammation was evaluated by synovitis score after 2-month treatment (n = 8). **f,** The subchondral bone plate (SBP) thickness of sham- or DMM-operated knee joints with 4-month treatment was revealed by representative 3D color maps. Color ranges from 0 (blue) to 320 μm (red). **g,** Quantitative analysis of the SBP thickness at the posterior site of femoral medial condyle after 4-month treatment (n = 8). **h,** Osteophytes on the sham- or DMM-operated knee joints with 4-month treatment was revealed by representative 2D (top) and 3D (bottom) microCT images. Red arrows indicate the osteophytes. **i,** Quantitative analysis of the total osteophyte volume after 4-month treatment (n = 8). **j,** von Frey assay on sham- or DMM-operated knee joints with 4-month treatment was performed at 1, 2, 4, 8, 12 weeks post-surgery (n = 8). The data of day 0 was acquired before DMM surgery. **k,** Representative images of immunohistochemistry staining of sPLA_2_-IIa, p-P65 and p-P100 in the tibial articular cartilage from sham- or DMM-operated knee joints with 2-month treatment. Scale bar, 100 μm. **l,** Quantification of sPLA_2_-IIa-, p-P65- and p-P100-positive chondrocytes in the tibial articular cartilage after 2-month treatment (n = 5). Statistical analysis was performed using oneway ANOVA with Turkey’s post hoc test. Data presented as mean ± s.e.m. *p<0.05, **p<0.01, ***p<0.001 in **c**, **e**, **g**, **i** and **l**. *p<0.05, **p<0.01 for sPLA_2_i-NPs vs. PBS in **j**.

The thickness of synovium is an indicator of inflammation in the knee joint (*33, 34*). PBS-, sPLA_2_i-, and Ctrl-NP-treated DMM knees exhibited a significantly thickened synovial lining with more than 10-fold increase in the synovitis score compared with sham knees. In contrast, sPLA_2_i-NP-treated DMM knees had only a 4-fold increase (Fig. 5d,e), suggesting that sPLA_2_i-NPs indeed generate an anti-inflammatory effect in the injured knees.

We further characterized some of the late OA symptoms including subchondral bone plate (SBP) sclerosis and osteophyte formation. At 4 months after DMM surgery, the thickness of the SBP at the femoral medial site in the knee joint was significantly increased in PBS-(1.25-fold), sPLA_2_i-(1.31-fold), and Ctrl-NP-(1.29-fold) treated groups compared to that in sham joints (Fig. 5f,g). However, this SBP sclerosis was not observed in the sPLA_2_i-NP-treated group. Furthermore, osteophytes were also frequently found in DMM knees in control treatment groups but not in the sPLA_2_i-NP-treated group (Fig. 5h,i). These data clearly indicate that sPLA_2_i-NPs prevent OA progression into a late stage.

In *WT* mice, mechanical allodynia was observed in the operated hind paw after DMM surgery but not following sham surgery (*35*). The mechanical threshold in DMM knees was greatly reduced compared to sham knees. Administration of PBS, sPLA_2_i, or Ctrl-NPs into DMM knees did not alter the OA pain (Fig. 5j). DMM knees receiving sPLA_2_i-NPs initially exhibited pain response at 1 week, probably due to the persistent post-surgical pain. At 2 weeks post-surgery, sPLA_2_i-NP treatment greatly increased the mechanical threshold to a level close to that in sham knees and maintained that level throughout the experimental period, implying that sPLA_2_i-NP treatment relieves OA pain.

We further studied the mechanism of sPLA_2_ inhibition for treating OA. IHC showed that sPLA_2_i-NPs blocked the up-regulation of sPLA_2_ after OA induction (Fig. 5k,l). Consequently, while DMM elevated the inflammatory signaling pathway, such as phosphor (p)-p65 and p-p100, sPLA_2_i-NP treatment greatly decreased the amounts of these inflammatory indicators in the cartilage at 2 and 4 months after DMM, thus protecting cartilage from OA insults (Fig. 5k,l and Supplementary Fig. S10). Collectively, these results suggest that the sPLA_2_ enzyme mediates the pathogenesis of OA, while sPLA_2_i-NPs inhibit OA inflammation in DMM knee joints.

To assess the toxicity of sPLA_2_i-NPs in mice, the histopathology of tissues from the liver, spleen, kidney, lung, heart and knee joint of mice was evaluated two months after serial intra-articular injections of PBS, sPLA_2_i, Ctrl-NPs or sPLA_2_i-NPs. We did not observe any histopathological abnormalities in any of the mouse organs and knee joints which received these injections directly (Supplementary Fig. S11,12). Furthermore, we evaluated toxic effects on blood parameters of mice following treatment and did not observe any clinical signs of toxicity (Supplementary Fig. S13). These results suggest that sPLA_2_i-NPs at the adopted dosages show no toxicity in mice.

### sPLA_2_i-NPs block joint damage in a single load-induced mouse PTOA model

The second mouse OA model we tested is a non-invasive mechanical loading model, which is clinically relevant to post-traumatic OA (PTOA) (Fig. 6a). This injury model allows for the study of early events after impact, at a time when inflammation is thought to be particularly important (*29*). We applied a single loading episode, composed of 60 cycles of 6 N or 9 N peak load, on the mouse tibia to induce joint translation and anterior cruciate ligament (ACL) rupture. Two weeks later, a lesion was seen alongside with proteoglycan loss in the lateral femoral articular cartilage surface (Fig. 6b). Quantification of the length of cartilage injury revealed significant damage areas in PBS-, sPLA_2_i-, and Ctrl-NP-treated knees (Fig. 6c and Supplementary Fig. S14). As expected, 9 N generated more severe damage than 6 N in all groups. In this model, chondrocyte apoptosis at the loading site contributes significantly to OA development. Interestingly, TUNEL staining revealed that sPLA_2_i-NP treatment greatly reduced the number of apoptotic chondrocytes in cartilage following loading (Fig. 6d,e and Supplementary Fig. S15). Examining the inflammatory pathway again revealed a reduction of p-P65 and p-P100 in sPLA_2_i-NP-treated knee joints, but not in sPLA_2_i or Ctrl-NP groups (Fig. 6f,g and Supplementary Fig. S16). Similar to the above DMM model, we found that sPLA_2_i-NPs suppressed the amount of sPLA_2_-IIA enzyme in cartilage. Synovitis scores suggested significant synovitis in PBS-, sPLA_2_i-, and Ctrl-NP-treated knees (Fig. 6h,i and Supplementary Fig. S17). A von Frey assay clearly showed that mice in these groups experienced OA pain (Fig. 6j and Supplementary Fig. S18). In contrast, injections of sPLA_2_i-NPs into knee joints immediately and 48 hours after loading remarkably reversed these PTOA symptoms regardless of loading force. Specifically, compared to PBS-treated knees, the length of cartilage injury in sPLA_2_i-NP-treated knees was decreased by 67% (6N) and 59% (9 N); synovitis score was reduced by 71% (6N) and 56% (9 N); and pain threshold was increased to a level similar to sham knees. Hence, our data demonstrate a joint protective role of sPLA_2_i-NPs in load-induced OA.

**Fig. 6.**
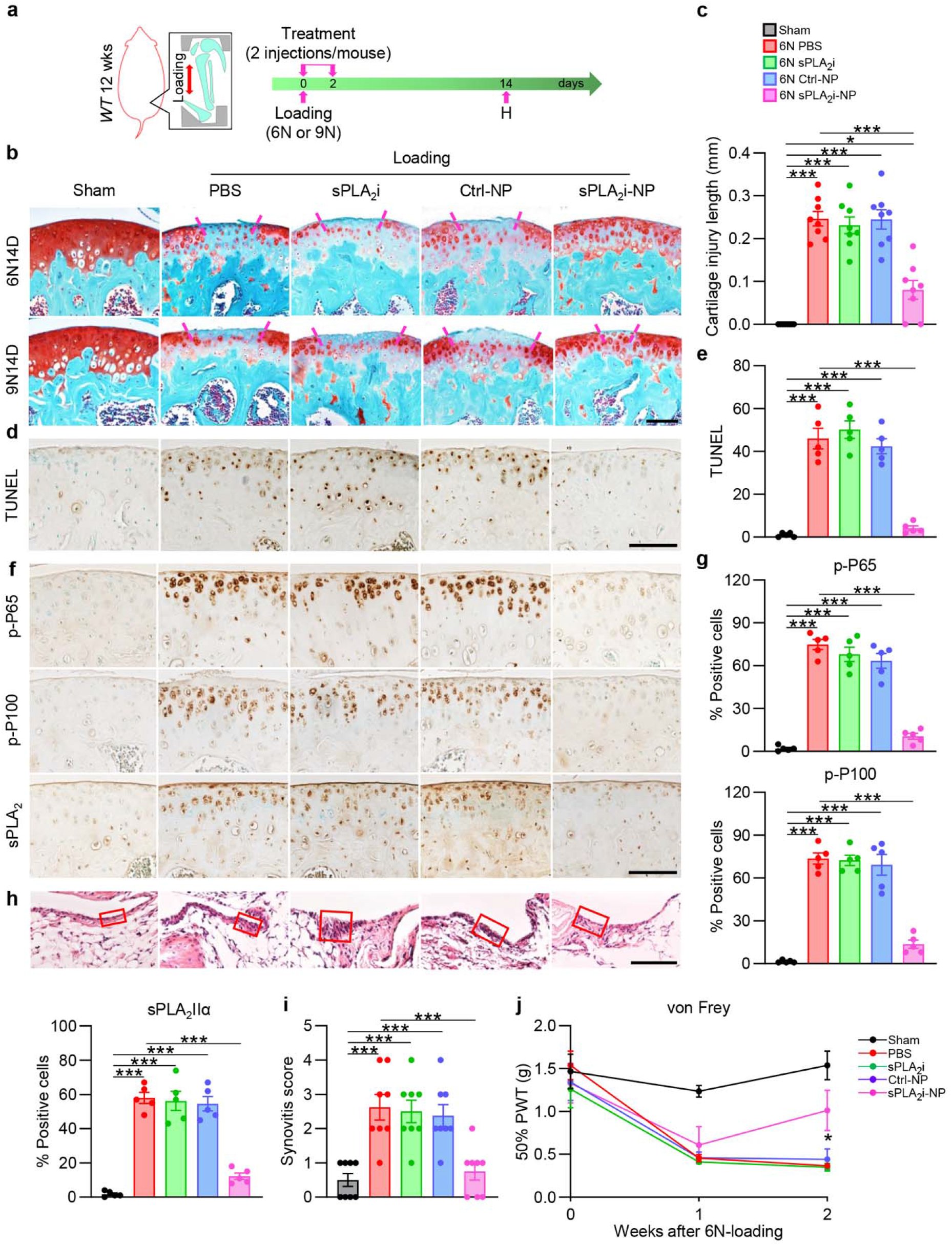
Therapeutic effects of sPLA_2_i-NPs for attenuation of load-induced post-traumatic OA. **a,** The study design of sPLA_2_i-NP treatment for loading-induced OA mice. 6N- or 9N-loading was performed on 3-month-old male mice. Intra-articular injections of PBS, sPLA_2_i, Ctrl-NPs and sPLA_2_i-NPs were made immediately and 48 hours post loading. **b,** Representative images of safranin O/fast green staining in the articular cartilage of sham- or load (6N or 9N)-operated knee joints with 14-day treatment. Scale bar, 100 μm. Dashed line indicate the range of loss of staining. **c,** Quantitative analysis of the length of the cartilage lesion range in the sham- or 6N-load-operated knee joints with 14-day treatment. **d,** Representative images of TUNEL staining in the tibial articular cartilage from sham- and 6N-load-operated knee joints with 14-day treatment. Scale bar, 100 μm. **e,** Quantification of TUNEL-positive chondrocytes in the tibial articular cartilage after 14-day treatment (n = 5). **f,** Representative images of immunohistochemistry staining of sPLA_2_-IIa, p-P65 and p-P100 in the tibial articular cartilage from sham- and 6N-load-operated knee joints with 14-day treatment. Scale bar, 100 μm. **g,** Quantification of sPLA_2_-IIa-, p-P65- and p-P100-positive chondrocytes in the tibial articular cartilage after 14-day treatment (n = 5). **h,** Representative images of H&E staining in the sham- and 6N-load-operated knee joints with 14-day treatment. Scale bar, 100 μm. Red boxed area indicate the enlargement of synovial lining cell layer. **i,** Synovial inflammation was evaluated by synovitis score after 14-day treatment (n = 8). **j**, von Frey assay on sham and 6N-load-operated knee joints with 14-day treatment at 1 or 2 weeks post loading (n = 8). The data of day 0 was acquired before loading. Statistical analysis was performed using one-way ANOVA with Turkey’s post hoc test. Data presented as mean ± s.e.m. *p<0.05, ***p<0.001 in **c**, **e**, **g** and **i**. *p<0.05 for sPLA_2_i-NPs vs. PBS in **j**.

## DISCUSSION

Growing evidence suggests that inflammation has a critical role in the pathogenesis of OA. Classical anti-inflammatory agents have limited utility to slow the progression of OA. In this work, we show that the level of sPLA_2_ is significantly increased in articular cartilage in both human and mouse OA knee tissues. Since the release of free AA from membrane phospholipids by sPLA_2_ is one of the major contributors to producing potent inflammatory mediators such as eicosanoids or platelet-activating factor, we therefore hypothesized that sPLA_2_ could act as a novel therapeutic target for OA treatment. Our strategy was to develop a sPLA_2_i-based approach for inhibition sPLA_2_ activity. Compared to classical anti-inflammatory agents, inhibiting sPLA_2_ enzyme activity by sPLA_2_i could offer the potential to block production of a more complete set of inflammatory substances through targeting upstream inflammatory pathways, leading to a safer and more effective alternative to conventional anti-inflammatory agents.

Great efforts from pharmaceutical companies and academic labs have recently been devoted to developing potent and selective sPLA_2_i for the treatment of inflammation-related diseases, with several ongoing clinical trials (*36*). One clinical trial used the sPLA_2_i LY315920 to treat patients with rheumatoid arthritis (RA). There was a significant short-term improvement after several days of treatment; however, no significant longer-term effect was observed after weeks of treatment (*37*). This failure is largely due to the lack of sufficient inhibitor concentration in the joint (*37, 38*), because small sPLA_2_i molecules have a high rate of clearance from the joint space. Our results also supported this limitation (i.e. fast clearance of sPLA_2_i from the joint), showing that intra-articular injection of free sPLA_2_i alone did not have a protective effect on cartilage degeneration in OA animals.

Drug delivery systems utilizing nanoparticles are increasingly being used to improve therapeutic delivery due to the favorable pharmacokinetics, biodistribution, and solubility of nanoparticles compared with free drugs. Currently, several nanoparticles have been developed and tested in pre-clinical OA studies, including liposomes, dendrimers, and poly(lactic-co-glycolic acid) (PLGA) particles. In this study, we have addressed the sPLA_2_i delivery challenges by incorporating small sPLA_2_i into nanometer-sized phospholipid micelles. By tuning the surface charge, the sPLA_2_i-NPs showed deep cartilage penetration and prolonged residence time in knee joints. This is especially important since the therapeutic target sPLA_2_ is located within deep cartilage tissue.

It should also be noted that most nanoparticles developed and tested in pre-clinical OA studies so far have some major weaknesses. Due to their relatively large size (100-200 nm in diameter), liposomes have prolonged residence times in joints, but are inaccessible to deep cartilage tissues (*39*), which could become problematic if therapeutic targets (i.e. sPLA_2_, the therapeutic target in this study) are located within deep cartilage tissue. Therefore, liposomes are only particularly well suited for OA drug delivery to the cartilage surface (*39, 40*). Due to its small size and controllable surface charge, dendrimers (i.e. PAMAM) can penetrate cartilage efficiently (*1*). However, a major weakness of dendrimers is the inability to encapsulate small sPLA_2_i by conjugation or physical interaction. In addition, dendrimers have not translated into the clinic despite 40 years of research (*41*). Poly (lactic-co-glycolic acid) (PLGA)-based particles is one of the most effective biodegradable polymeric particles and the only FDA-approved IA delivery system (*42*). However, due to their negative surface charge (*43*), the PLGA-based drug delivery systems are not effective for targeting middle and deep cartilage unless extremely high drug doses are used (*39, 44*). Compared to the above nanoparticles, phospholipid micelles we used here for loading sPLA_2_i have several advantages. First, sPLA_2_i-NPs consist almost solely of clinically used phospholipids, and thus are readily translatable to clinical applications. Second, lipid-based sPLA_2_ inhibitors can be incorporated into phospholipid micelles with high yield and without any conjugation or purification steps. Third, sPLA_2_i-NPs allow sustained and controlled drug release based on the pathological conditions. Fourth, the method for producing sPLA_2_i-NPs is highly reproducible, simple and cost-effective since all components are included in starting materials prior to sPLA_2_i-NP formation, which would allow large-scale, GMP production of nanoparticles, a necessary step for the initiation of future clinical trials.

Inflammation plays a critical role in OA progression. Compared to rheumatoid RA, the prototypical inflammatory arthritis, the inflammation in OA is chronic and relatively low-grade (*45*). Systemically blocking the activity of conventional inflammatory cytokines with biologic therapies approved for the treatment of RA, such as anti-TNF or anti-IL-1β therapies, provides no or minor benefit in generalized OA (*45*). These observations implicate that a new target or approach must be adopted for treating OA inflammation and resultant cartilage degeneration. Inflammation starts with the PLA_2_-induced hydrolysis of the membrane phospholipids, giving rise to the fatty acids, such as AA and lysophospholipids. Of all PLA_2_ types, sPLA_2_ is considered the most “inflammatory enzyme”. Past studies found that sPLA_2_ amount increases in human OA joints, including cartilage chondrocytes (*20–23*), but they did not further analyze the cause-effect relationship between sPLA_2_ upregulation and OA progression. Our studies first confirmed the enhanced sPLA_2_ activity in both human and mouse OA cartilage. Then we used an ex vivo bovine explant model to demonstrate that inhibiting sPLA_2_ protects chondrocytes against IL-1β insult via suppressing inflammation pathways. Most importantly, we used two in vivo mouse models of OA, which mimic different OA subtypes, to provide proof-of-principle evidence that intraarticular injections of sPLA_2_i-NPs are effective in attenuated OA progression. Strikingly, local delivery of sPLA_2_i-NPs, but not free sPLA_2_i nor nanoparticle alone, greatly slowed the progression of cartilage degeneration, reduced synovial inflammation, prevented osteophyte formation, and relieved join pain. Meanwhile, no signs of toxicity have been observed in the joints and other tissues, suggesting the safety of this nanomedicine. That to be said, since OA is a chronic disease, long term treatment is required in the future to further prove its safety.

The future development of sPLA_2_i-NPs-based therapies requires further optimization and investigation. For example, in this work we used thioetheramide-PC as sPLA_2_i because of its commercial availability and its ease of loading into phospholipid micelles. However, thioetheramide-PC is for research use only and has a relatively high half-maximal inhibitory concentration (IC_50_ ~2 μM). In the future, we will choose sPLA_2_ inhibitors that have already been used in clinical trials and have a low IC_50_ value. There are some other variables that can be optimized to further improve the therapeutic efficacy of sPLA_2_i-NPs, including particle size, surface charge, and surface-conjugated ligands. It should be mentioned that the sPLA_2_i-NPs administration was initiated immediately after DMM surgery in this study. Future animal studies should incorporate injections at time points when early or middle-stage OA symptoms appear. Given that sPLA_2_ level was drastically elevated in human knee cartilage at early-, middle-, and late-stage of OA progression, it is reasonable to expect that the sPLA_2_i-NPs could also be used to slow cartilage degeneration and OA progression in cases where OA is already established. In addition, preclinical investigation in larger animals is necessary for clinical translation and regulatory approval.

## MATERIALS AND METHODS

### Materials

1-Palmitylthio-2-palmitoylamido-1,2-dideoxy-*sn*-glycero-3-phosphorylcholine (thioetheramide-PC) was purchased from Cayman Chemical (Ann Arbor, Michigan). Hydrogenated soy phosphatidylcholine (HSPC), 1,2-distearoyl-*sn*-glycero-3-phosphoethanolamine-N-[methoxy(polyethylene glycol)-2000] (DSPE-PEG2000), 1,2-dioleoyl-3-trimethylammonium-propane (DOTAP), 1,2-dioleoyl-sn-glycero-3-phosphoethanolamine-N-(Lissamine rhodamine B sulfonyl) (Rhod-PE), 1,2-distearoyl-sn-glycero-3-phosphoethanolamine-N-[amino(polyethylene glycol)-2000]-N-(Cyanine 7) (DSPE-PEG2000-Cy7), and1-palmitoyl-2-{6-[(7-nitro-2-1,3-benzoxadiazol-4-yl)amino]hexanoyl}-*sn*-glycero-3-phosphocholine (NBD-PC) were purchased from Avanti Polar Lipids, Inc. Secreted phospholipase A_2_ (PLA_2_) enzyme from Naja mossambica was purchased from Sigma-Aldrich Co. Rabbit polyclonal antibody to sPLA_2_-IIa, phospho-P65 (p-P65) and phospho-P100 (p-P100) antibodies were purchased from Abcam. All other chemicals were used as received. All of the buffer solutions were prepared with deionized water.

### Synthesis of sPLA_2_i-loaded phospholipid nanoparticles

sPLA_2_i-NPs were prepared by hydration of dry sPLA_2_i/lipid films. A mixture containing 10□mol% DOTAP/25□mol% thioetheramide-PC /65□mol% DSPE-PEG2000 was prepared in a round bottom flask. The total amount of thioetheramide-PC was 0.25 mg. For the preparation of fluorescently labeled nanoparticles, a small amount (1 mol%) of the fluorescent lipids, Rhod-PE or DSPE-PEG2K-Cy7, was also added to the thioetheramide-PC /lipid mixture. The solvent was removed using a direct stream of nitrogen prior to vacuum desiccation for a minimum of 4 hours. Nanoparticles were formed by adding an aqueous solution (0.1 × PBS, pH = 7.4) to the dried film and incubating in a 25 °C water bath for 5 minutes and then votexing for another 3 minutes. The resulting solution was then centrifuged at 3000 g for 5 minutes to remove the large aggregates.

Finally, the nanoparticles were filtered through a 0.22 μm cellulose acetate membrane filter (Nalgene, Thermo Scientific) and stored in the dark at 4 °C. Control nanoparticles, including nanoparticles without sPLA_2_i (10□mol% DOTAP/25□mol% HSPC/65□mol% DSPE-PEG2000) and nanoparticles without DOTAP (10□mol% HSPC/25□mol% sPLA_2_i/65□%mol% DSPE-PEG2000), were prepared using the similar procedures as above.

Synthesized sPLA_2_i-NPs were incubated in water at 4 °C as well as in bovine synovial fluid at 37 °C for the stability study. The diameter and zeta potential of the nanoparticles were measured with dynamic light scattering (DLS, Malvern, Zetasizer, Nano-ZS). The morphology of the nanoparticles was observed using a transmission electron microscope (TEM) (JOEL 1010) by the negative-staining technique.

### In vitro sPLA_2_ response study

To study in vitro sPLA_2_ response of sPLA_2_i-NPs, the NBD-incorporated liposomes were incubated with sPLA_2_i-NPs in the presence of sPLA_2_ enzyme and the dequenching of NBD fluorescence signal was monitored. To prepare NBD-liposomes, stock solutions of HSPC and NBD-PC in chloroform were mixed in the following molar ratios: HSPC/NBD-PC (80:20). The total amount of HSPC was 1 mg. The solvent was removed using a direct stream of nitrogen prior to vacuum desiccation for a minimum of 4 hours. 0.2 ml deionized water was then added to the dried lipid film and incubated in a 50 °C water bath for 0.5 hours and then sonicated for another 30 minutes. The stock solution of NBD-incorporated phospholipid liposomes was stored in the dark at 4 °C.

Dequenching measurements were performed by first preincubating a mixture of either sPLA_2_i-NPs or Ctrl-NPs (without sPLA_2_i) with the sPLA_2_ enzyme (10 μL sPLA_2_i-NPs [sPLA_2_i: 0.25 mg ml^-1^] or Ctrl-NPs + 6.67 μL sPLA_2_ enzyme [sPLA_2_: 7.5 U ml^-1^]) for 20 minutes. Fluorescence measurements of the NBD-incorporated liposomes in buffer (20 μL of NBD-liposomal suspension ([HSPC] = 1 mg ml^-1^) + 0.48 ml of 10 mM HEPES (pH 7.4) buffer solution containing 2 mM CaCl_2_) were taken for 5 minutes prior to the addition of the incubation mixture of the sPLA_2_i-NPs or Ctrl-NPs. The fluorescence intensity at 520 nm was measured on a SPEX FluoroMax-3 spectrofluorometer (Horiba Jobin Yvon) using an excitation at 460 nm. The amount of NBD dequenched (% NBD dequenched) was calculated by means of Equation 1:

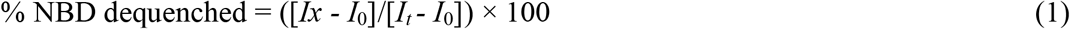

where *I*_0_ is the fluorescence intensity of the liposomal suspension containing NBD at the initial time, *Ix* is the fluorescence intensity at any given time, and *I*_t_ is the fluorescence intensity after addition of 20 μL Triton X-100 (50 mM) to the suspension at the end of experiment.

### Cell culture

Primary mouse chondrocytes were isolated from the distal femoral and proximal tibial epiphysis of mice (3-6 days old) via enzymatic digestion. Briefly, tissues were incubated with 0.25% trypsin (Invitrogen) for 1 h, followed by 2 h digestion with 300 U ml^-1^ type I collagenase (Worthington Biochemical). Dissociated cells were seeded in culture plates, and attached cells were considered primary mouse chondrocytes. They were cultured in DMEM/F12 containing 10% FBS, 100 μg ml^-1^ streptomycin, and 100 U ml^-1^ penicillin. Primary mouse chondrocytes between passage 0 and passage 3 were used for MTT assay experiments.

### MTT assay

Primary mouse chondrocytes (5000 cells per well) were seeded in 96-well plates and incubated overnight (37 °C, 5% CO_2_) to allow the cells to attach to the surface of the wells. The sPLA_2_i-NPs were added to wells at five different sPLA_2_i concentrations ranging from 0.625 to 10 μg ml^-1^ (0.625, 1.25, 2.5, 5, 10 μg ml^-1^), and the cell viabilities were determined according to the supplier’s instructions. After 24 hours incubation, the medium containing nanoparticles in each well was aspirated off and replaced with 100 μL fresh medium and 10 μL of MTT reagent. The cells were incubated for 2 to 4 hours. Then, 100 μL detergent reagent was added and left at room temperature in the dark for 4 hours. Finally, the absorbance of formazan product was measured on a Tecan microplate reader (BioTek Instruments, Inc) at 570 nm. Cell viability was calculated using the following equation:

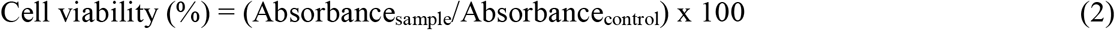

### Bovine cartilage explant harvest and culture

Young (1-2 weeks old) bovine knee joints were obtained from Vendors (Lampire biological laboratories), and cartilage explants were harvested from the trochlear groove using biopsy punch and cultured with chemically defined medium (DMEM, 1% ITS + Premix, 50 μg ml^-1^ L-proline, 0.1 μM dexamethasone, 0.9 mM sodium pyruvate and 50 μg ml^-1^ ascorbate 2-phosphate) in 48-well plate.

### Bovine cartilage explant penetration assay

Bovine cartilage explants (6 *mm* in *diameter* and *2 mm* in *thickness*) were incubated with free rhodamine, rhodamine-labeled sPLA_2_i-NPs (DOTAP-) or sPLA_2_i-NPs (DOTAP+) in 500 μl of culture medium for 48, 96, 144 or 192 hours at 37 □ and 5% CO_2_ under gentle agitation. Culture medium with free rhodamine, rhodamine-labeled sPLA_2_i-NPs (DOTAP-) or sPLA_2_i-NPs (DOTAP+) was replaced every other day. In all cases, the final Rhodamine concentration in the culture medium was 10 μM. After incubation, cartilage explants were washed three times with PBS, fixed with 4% Paraformaldehyde (PFA), dehydrated with 20% sucrose + 2% PVP (Polyvinylpyrrolidone) followed by embedding with 30% sucrose + 2% PVP + 8% Gelatin. Sections were mounted with DAPI Fluoromount-G Mounting Medium on glass slides and immediately observed under confocal microscope (Zeiss LSM 710). Quantitative analysis was performed on maximum intensity projections of Z-stack images taken from 100 μm thick sections.

### Bovine cartilage explant uptake assay

A total volume of 300 μL of free rhodamine, rhodamine-labeled sPLA_2_i-NPs (DOTAP-) or sPLA_2_i-NPs (DOTAP+) in culture medium was added to bovine cartilage explants (3 *mm* in *diameter* and *2 mm* in *thickness*). The final rhodamine concentration in the culture medium was 10 μM. The explants were incubated for 24 hours at 37 □ and 5% CO_2_ under gentle agitation. The explants were then removed from the medium, washed tree times with PBS, imaged by IVIS (Spectrum, PerkinElmer). Radiant efficiency within a fixed anatomical region of interest (ROI) was measured using Living Image software.

### In vivo joint retention assay

The mouse knee joints retention assay was assessed by intra-articular injection of 10 μl of 10 μM free ICG or 10 μM Cy7 doped sPLA_2_i-NPs in healthy (3 months old) and OA (8 weeks post DMM surgery) mouse knees. The rat knee joints retention assay was assessed by intra-articular injection of 40 μl of 10 μM free ICG or 10 μM Cy7 doped sPLA_2_i-NPs in healthy rat knees (3 months old). An IVIS (Spectrum, PerkinElmer) was used to serially acquire fluorescence images within each joint over a period of 4 weeks. Using Living Image software, radiant efficiency within a fixed anatomical region of interest (ROI) was measured.

### In vivo biodistribution assay

In vivo biodistribution study was performed by intra-articular injection of 10 μl of PBS or 10 μM Cy7 doped sPLA_2_i-NPs in mouse knees (3 months old). At 24 hours or 1 month following injection, the mice were sacrificed. The knee joints, blood, and major organs (heart, liver, spleen, lung, kidney) were harvested. Knees were dissected to isolate the major joint components, including the surrounding tissues (quadriceps, patella, patellar ligament, synovium, fat pad), femoral condyles, tibial plateau and meniscus. All the major joint components, blood and organs were imaged using the IVIS, and the data was analyzed as described above.

### Mouse femoral head explants penetration assay

Mouse femoral heads were collected from 8-week-old male mice and cultured for 48□hours in chemically defined medium in 48-well plate. Following the culture, mouse femoral head explants were stimulated using 10□ng□ml^-1^ recombinant mouse IL-1β (PeproTech) for 2 days. On day 3, rhodamine-labeled sPLA_2_i-NPs were incubated with the IL-1β-stimulated femoral heads for 24 or 48 hours at 37□°C with gentle agitation. The final rhodamine concentration in the culture medium was 10 μM. Following incubation, femoral heads were washed with PBS, fixed with 4% PFA, dehydrated with 30% sucrose and embedded in Tissue-Tek OCT Compound. 6-μm-thick cyrosections were cut and mounted with DAPI Fluoromount-G Mounting Medium on the slides and imaged with a fluorescence microscope (Nikon, Eclipse 90i). Images were analyzed with ImageJ to quantify the penetration depth nanoparticle into the cartilage. Fluorescence intensity within each image was measured with ImageJ and normalized to the fluorescence of the outermost cartilage surface of the treated femoral head.

### Mouse femoral head explants degradation assay

Mouse femoral heads were collected using the same procedure as described in mouse femoral head explants penetration assay. To test the therapeutic effects of sPLA_2_i-NPs, femoral head cartilage explants were divided into 4 groups to receive PBS (i.e. untreated), IL-1β, IL-1β and Ctrl-NP, and IL-1β and sPLA_2_i-NP treatments, respectively, for 8 days. The final IL-1β and sPLA_2_i concentration in the culture medium was 10□ng□ml^-1^ and 0.1 mg□ml^-1^, respectively. The culture medium was replaced at day 2, 4, and 6. After 8-day incubation, mouse femoral heads were then fixed with 4% paraformaldehyde overnight followed by decalcification in 0.5 M EDTA (pH = 7.4) for 2 weeks and processed for 6-μm paraffin sections. Paraffin sections were used for safranin-O staining, and the safranin-O-positive area was quantified with ImageJ. OA severity was evaluated by Mankin score. Both quantifications were based on the section with the most severe loss of safranin-O staining and cartilage damage.

### Quantitative RT-PCR

Mouse femoral heads were collected and treated using the same procedure in femoral head explants degradation assay. Total RNA was harvested from the femoral head articular cartilage using Tri Reagent (Sigma). Taqman Reverse Transcription kit (Applied BioSystems) was used to reverse-transcribe mRNA into cDNA. Following this, PCR was performed using a Power SYBR Green PCR MasterMixkit (Applied BioSystems). The primer sequences for the genes used in this study are listed in Table 1.

### Human articular cartilage samples

The human OA articular cartilage samples were prepared from the de-identified specimens obtained at the total arthroplasty of the knee joints and used for immunohistochemical examination for sPLA_2_. The serial sections were stained by Safranin O and Fast green staining to evaluate OA phenotype and stage.

### Animal care

In accordance with the standards for animal housing, mice were group housed in an atmosphere of 23-25°C with a 12-hour light/dark cycle, and allowed free access to water and standard laboratory pellets. All work performed on animals was approved by the Institutional Animal Care and Use Committee at the University of Pennsylvania.

To induce mouse OA that mimics chronic OA in human patients, male mice at 3 months of age were subjected to DMM surgery at right knees and sham surgery at left knees. Briefly, in DMM surgery, the joint capsule was opened immediately after anesthesia and the medial meniscotibial ligament was cut to destabilize the meniscus without damaging other tissues. In Sham surgery, the joint capsule was opened in the same fashion but without any further damage.

To induce mouse OA that is noninvasive and mimics post-traumatic OA in human patients, male mice (2 months old) were subjected mechanic loading at the right knees and sham loading at left knees. Briefly, under anesthesia, the right tibiae were positioned with the knee downward in deep flexion between custom-made cups and subjected to axial compressive loads with a peak force of 6 or 9 Newtons (N) with a 0.5 N preload force to maintain the limb in position between loading cycles. Cyclic loads were applied for 0.34 s with a rise and fall time each of 0.17 s and a baseline hold time of 10 s between cycles for 60 cycles. The uninjured left knees were used as controls.

### Nanoparticle administration

For treatment, nanoparticles were administrated using sterile techniques: the right knees were kept in a flexed position and a total volume of 10 μl of PBS, sPLA_2_i (0.25 mg ml^-1^), Ctrl-NPs, or sPLA_2_i-NPs (sPLA_2_i: 0.25 mg ml^-1^) was injected intra-articularly with a 30-gauge needle. For DMM surgical OA model, the first injection was performed *immediately* after surgery. Injections were then repeated every 2 weeks for 2 or 4 months. In total, there are 4 injections for the 2 months group and 8 injections for 4 months group. For load-induced OA model, injections (i.a.) were performed *immediately* and at 48 hours after loading. There are 2 injections in total for each mouse in this model.

After 2 months treatment for DMM mice, some major organs (kidney, liver, lung, heart, and spleen) and blood were collected. Tissue sections were stained with hematoxylin and eosin (H&E) to assess the effects of different treatment on mouse organ morphology. The blood indexes were measured after receiving 2-month treatment of PBS, sPLA_2_i, Ctrl-NPs, or sPLA_2_i-NPs.

### Histology

After euthanasia, mouse knee joints were harvested and fixed in 4% paraformaldehyde overnight followed by decalcification in 0.5 M EDTA (pH 7.4) for 4 weeks prior to paraffin embedding. A serial of 6 μm-thick sagittal sections (about 100) were cut across the entire medial compartment of the joint until ACL junction. To measure the thicknesses of articular cartilage and chondrocyte numbers, 3 sections from each knee, corresponding to 1/4 (sections 20-30), 2/4 (sections 45-55), and 3/4 (sections 70-80) regions of the entire section set, were stained with Safranin O/Fast green and quantified using BIOQUANT software. The final measurement is an average of these three sections. We defined uncalcified cartilage as the area from articular surface to the tide mark and calcified cartilage as the area from tide mark to cement line. The method to measure Mankin Score was described previously (*46*). Briefly, two sections within every consecutive six sections in the entire section set for each knee were stained with Safranin O/Fast green and scored by two blinded observers. Each knee received a single score representing the maximal score of its sections.

Synovitis score grading was carried out in 6-μm paraffin sections of sagittal mouse knee sections stained with H&E. The following basic morphological parameters of synovitis were included: (i) hyperplasia/enlargement of synovial lining layer, (ii) degree of inflammatory infiltration and (iii) activation of resident cells/synovial stroma, including fibroblasts, endothelial cells, histiocytes, macrophages, and multinucleated giant cells. All parameters are graded from 0 (absent), 1 (slight), 2 (moderate) to 3 (strong positive).

Cartilage injury length was measured in Safranin O/Fast green stained paraffin sections. From the serial Safranin-O stained sections in each sample, we selected one section with the widest cartilage lesion that featured by focal loss of Safranin-O staining, minor fissuring of articular cartilage, and atrophy of articular chondrocytes. According to these histological changes, it is possible to identify the demarcation between the normal and injured cartilage tissue, and the length of the cartilage lesion range was measured.

Paraffin sections were used for immunohistochemistry and TUNEL assay. For mouse samples, after appropriate antigen retrieval, slides were incubated with primary antibodies, such as rabbit anti-sPLA_2_-IIA (Abcam, ab23705), rabbit anti-p-P65 (Abcam, ab86299), rabbit anti-p-P100 (Abcam, ab194919) at 4 °C overnight, followed by binding with biotinylated secondary antibodies and DAB color development. The TUNEL assay was carried out according to the manufacturer’s instructions (Millipore, s7101). For human samples, anti-sPLA_2_-IIA (Abcam, ab23705) antibody was used.

### OA pain analysis

The knee joint pain after DMM surgery or loading injury was evaluated in mice weekly before and after surgery using von Frey filaments as described previously (*47*). An individual mouse was placed on a wire-mesh platform (Excellent Technology Co.) under a 4 × 3 × 7 cm cage to restrict their move. Mice were trained to be accustomed to this condition every day starting from 7 days before the test. During the test, a set of von Frey fibers (Stoelting Touch Test Sensory Evaluator Kit #2 to #9; ranging from 0.015 to 1.3 g force) were applied to the plantar surface of the hind paw until the fibers bowed, and then held for 3 seconds. The threshold force required to elicit withdrawal of the paw (median 50% withdrawal) was determined five times on each hind paw with sequential measurements separated by at least 5 minutes.

### Micro-computed tomography (microCT) analysis

The distal femur of mouse knee joints was scanned at a 6-μm isotropic voxel size with a microCT 35 scanner (Scanco Medical AG, Brüttisellen, Switzerland). All images were smoothened by a Gaussian filter (sigma = 1.2, support = 2.0). Measurement of SBP thickness was described previously (*48*). Briefly, sagittal images were contoured for the SBP followed by generating a 3D color map of thickness for the entire SBP along with a scale bar. This map was then converted to a grayscale thickness map. The region of interest (ROI) was circled and the average SBP thickness within ROI is calculated by average grey value/255 * max scale bar value. Coronal images were also contoured for the osteophyte followed by 3D reconstruction and volume calculation.

### Statistical analysis

Data are expressed as means ± standard error (s.e.m) and analyzed by t-tests, one-way ANOVA with Dunnett’s or Turkey’s posttest and two-way ANOVA with Bonferroni’s or Turkey’s posttest for multiple comparisons using Prism 8 software (GraphPad Software, San Diego, CA). For assays using primary chondrocytes and bovine cartilage explants, experiments were repeated independently at least three times and representative data were shown here. Values of p<0.05 were considered statistically significant.

## Acknowledgments

This study was supported by NIH grants NIH/NIAMS R01AR066098, R01DK095803, R21AR074570 (to L.Q.), P30AR069619 (to Penn Center for Musculoskeletal Disorders), R01NS100892 (to Z.C.).

