## supporting information for "Phospholipase A_2_ inhibitor-loaded micellar nanoparticles attenuate inflammation and mitigate osteoarthritis progression"

**This PDF file includes:**

Figure S1 to S18

Table S1

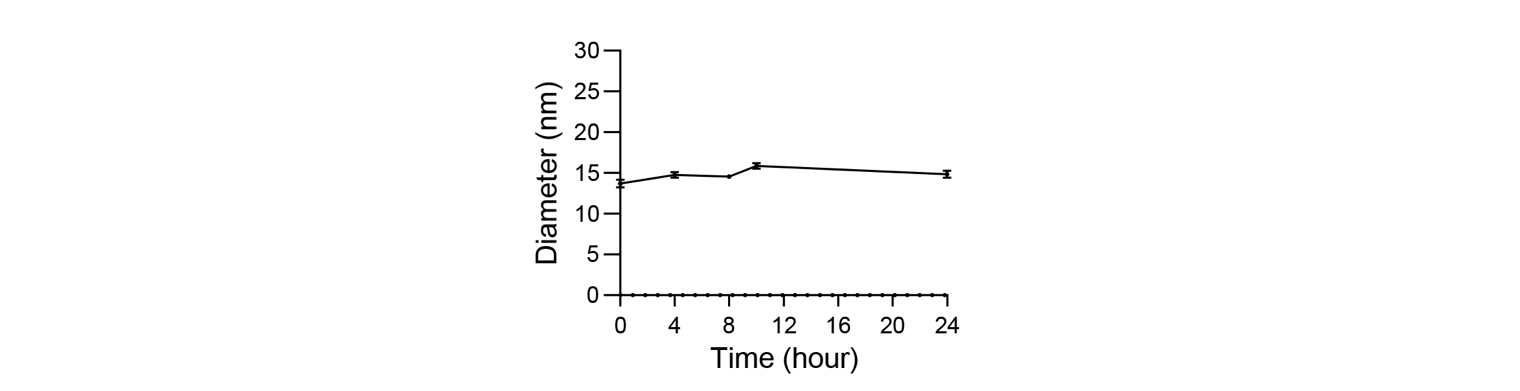

**Supplementary Fig. 1. The stability of sPLA2i-NPs in synovial fluid.** The stability of sPLA_2_i-NPs in synovial fluid was accessed by monitoring the hydrodynamic size (diameter) for up to 24 hours.

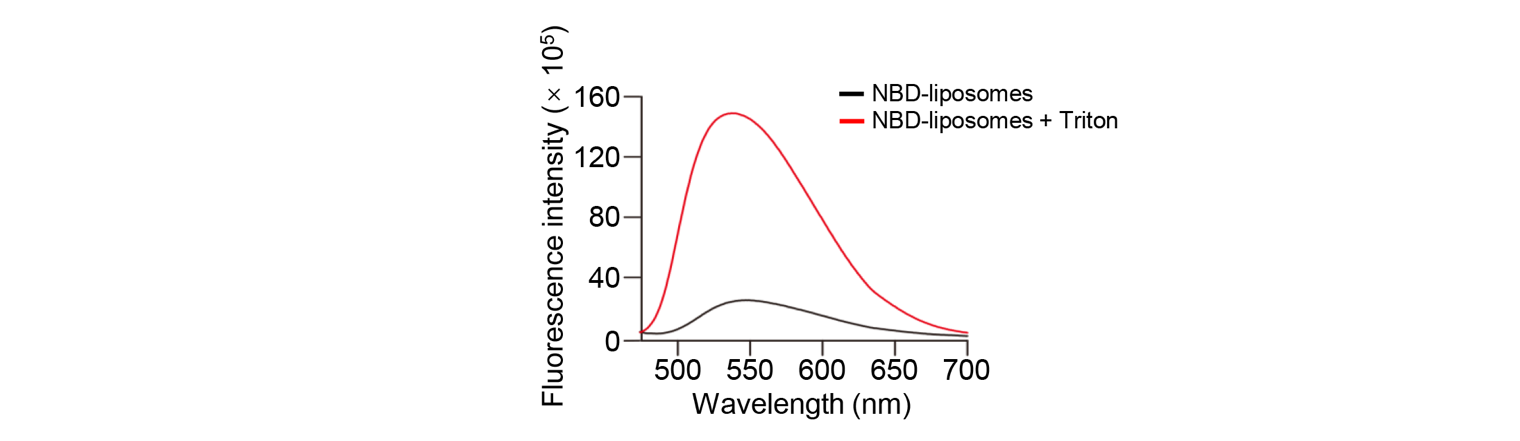

**Supplementary Fig. 2.** **Quenching study of NBD-liposomes (20 mol% NBD-PC/80 mol% HSPC liposome).**

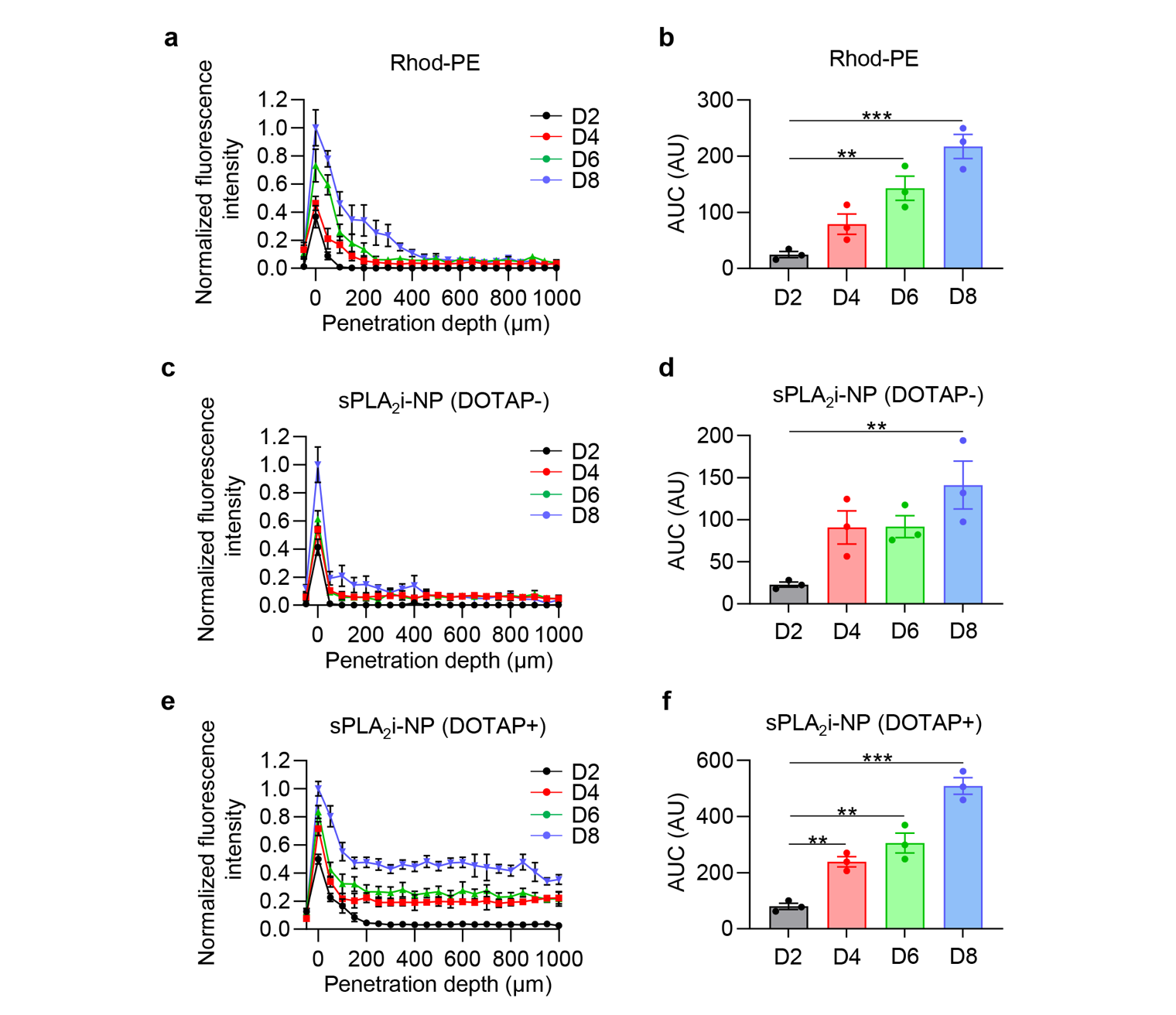

**Supplementary Fig. 3. Penetration ability of sPLA2i-NPs. a, c, e,** Quantitative analysis of free rhodamine dye (a), rhodamine-labeled sPLA_2_i-NPs (DOTAP-) (c), and sPLA_2_i-NPs (DOTAP+) (e) penetration depth into bovine cartilage explants over 8-day incubation (n = 3). **b, d, f,** Quantitative analysis of area under the curve (AUC) based on the corresponding fluorescence intensity profiles in a, c, and e, respectively (n = 3). Rhod-PE: free rhodamine dye, sPLA_2_i-NPs in the absence of DOTAP: sPLA_2_i-NPs (DOTAP-), sPLA_2_i-NPs in the presence of DOTAP: sPLA_2_i-NPs (DOTAP+). Statistical analysis was performed using one-way ANOVA with Dunnett’s post hoc test. Data presented as mean ± s.e.m. **p<0.01, ***p<0.001.

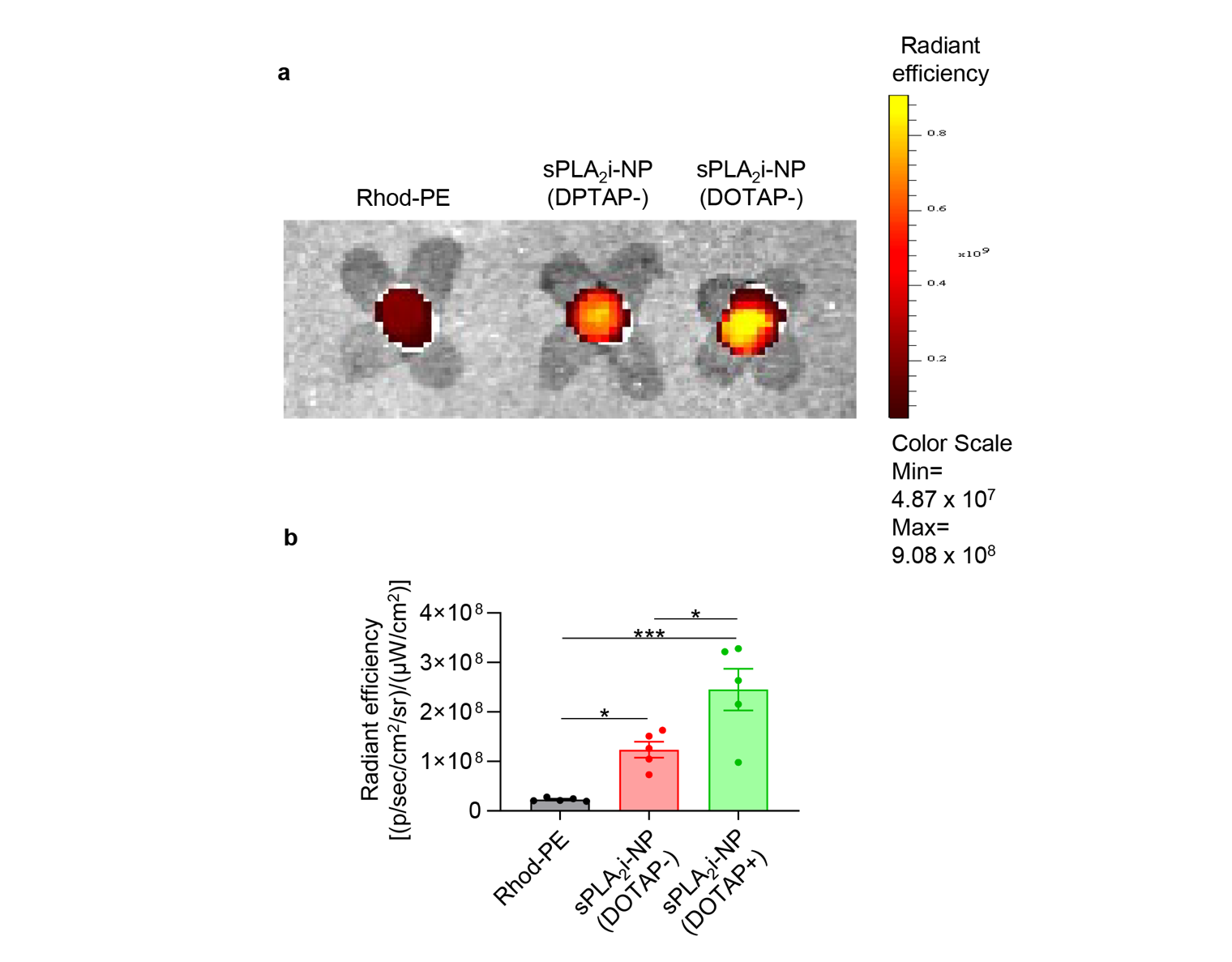

**Supplementary Fig. 4. sPLA_2_i-NP uptake in bovine cartilage explants. a,** Representative IVIS images of bovine cartilage explants incubated with free rhodamine dye, rhodamine-labeled sPLA_2_i-NPs (DOTAP-), and sPLA_2_i-NPs (DOTAP+) for 24 hours. Fluorescent scale, min = 4.87 × 10^7^, max = 9.08 × 10^8^. **b,** Quantitative analysis of fluorescent radiant efficiency on the surfaces of bovine cartilage explants after 24-hour incubation (n = 8). Rhod-PE: free rhodamine dye, sPLA_2_i-NPs in the absence of DOTAP: sPLA_2_i-NPs (DOTAP-), sPLA_2_i-NPs in the presence of DOTAP: sPLA_2_i-NPs (DOTAP+). Statistical analysis was performed using one-way ANOVA with Turkey’s post hoc test. Data presented as mean ± s.e.m. *p<0.05, ***p<0.001.

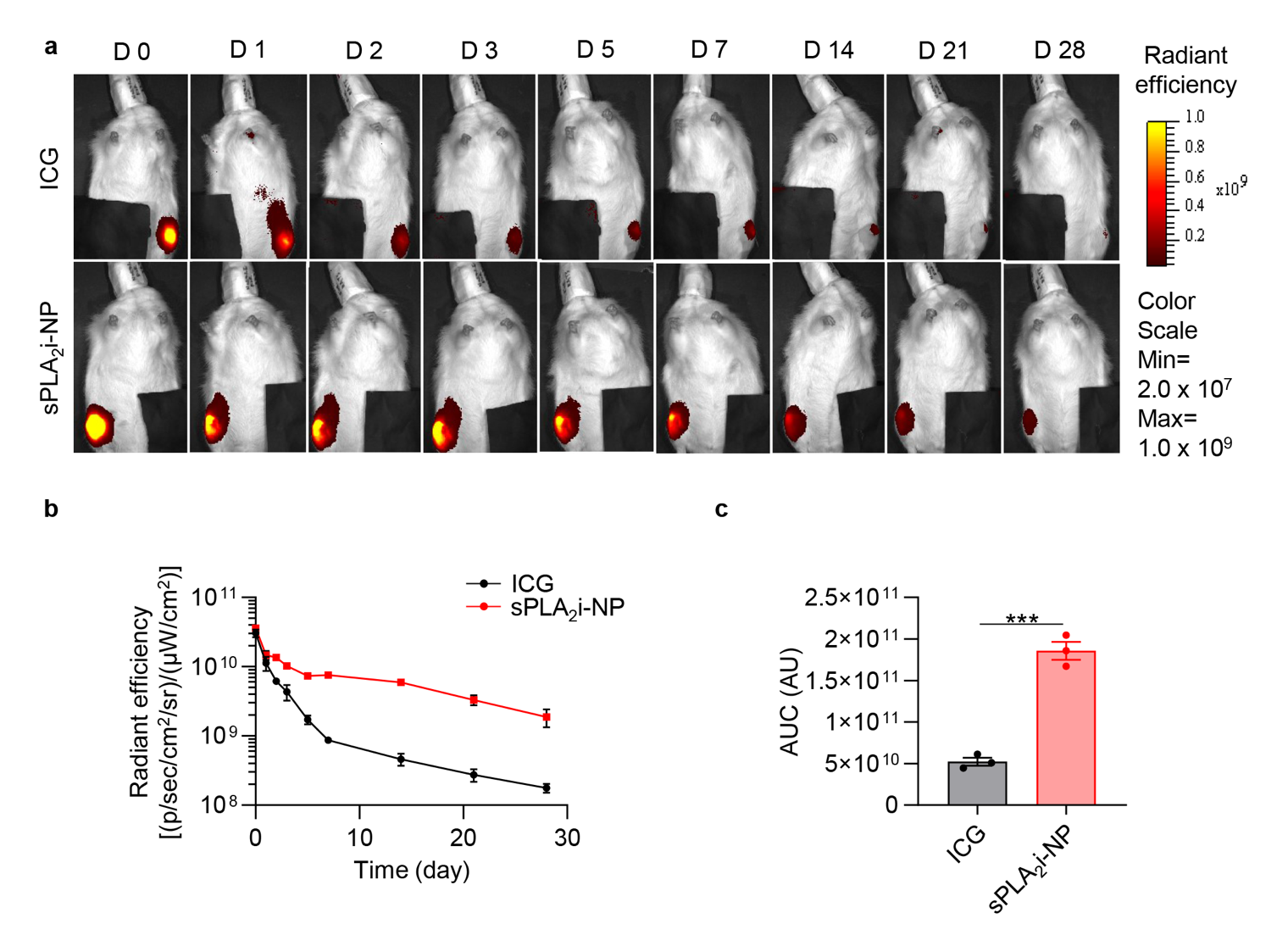

**Supplementary Fig. 5. The retention ability of sPLA_2_i-NPs in rat knee joints. a,** Representative IVIS images of healthy rat knee joints over 28 days post single intra-articular injection of Cy7-labeled sPLA_2_i-NPs or free ICG. Fluorescent scale, min = 4.0 × 10^7^, max = 1.0 × 10^9^. **b,** Quantitative analysis of time course fluorescent radiant efficiency within healthy rat knee joints over 28 days (n = 3). **c,** Quantification of the area under the curve (AUC) based on the fluorescence intensity profiles in b (n = 3). Statistical analysis was performed using paired two-tailed t-test. Data presented as mean ± s.e.m. ***p<0.001.

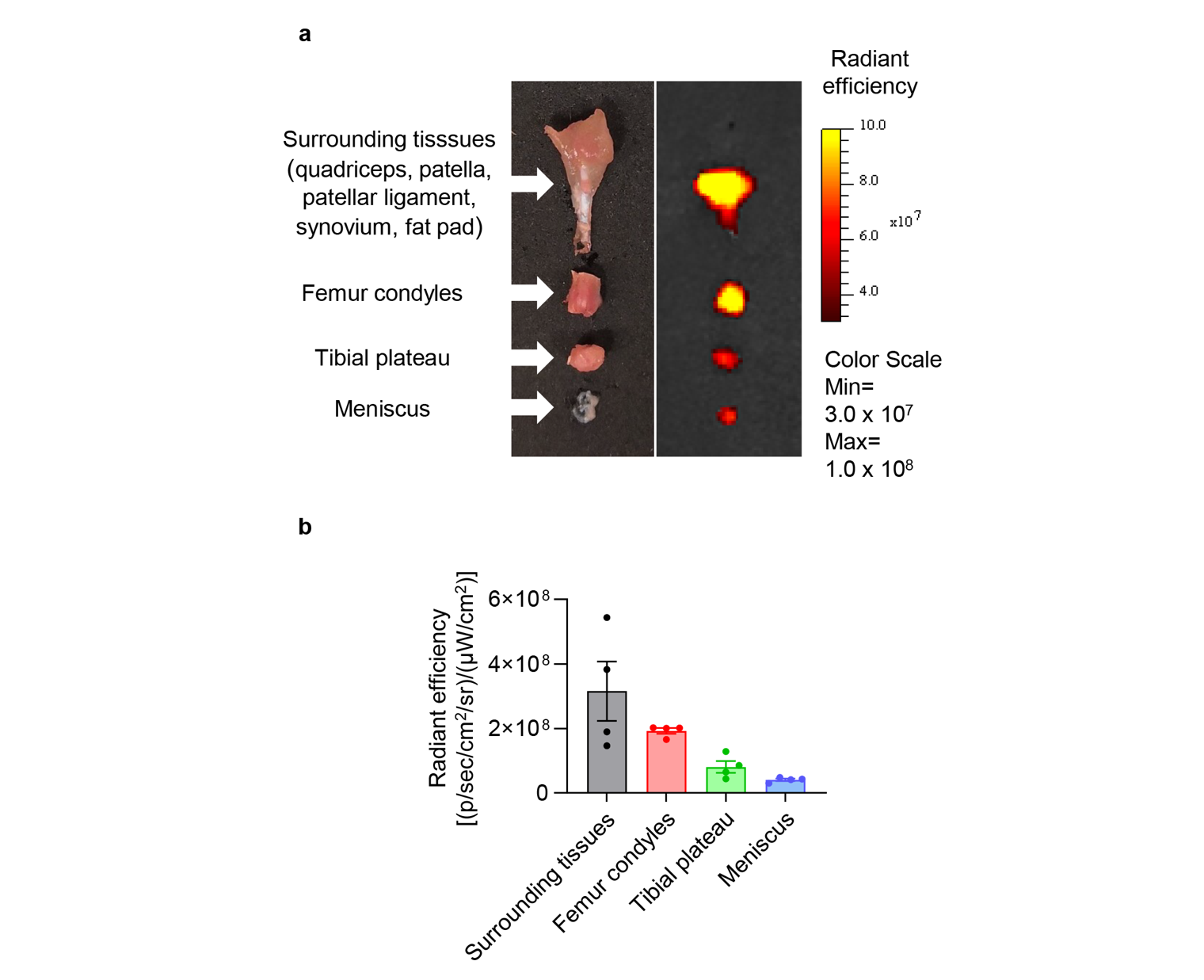

**Supplementary Fig. 6.** **Biodistribution of sPLA_2_i-NP within mice knee joint following local injection. a,** Biodistribution of Cy7-labeled sPLA_2_i-NPs within healthy mouse knee joints at 24 hours post single injection of Cy7-labeled sPLA_2_i-NPs. Fluorescent scale: min = 3.0 × 10^7^, max = 1.0 × 10^8^. **b,** Quantification of fluorescent radiant efficiency in the different components of knee joints (n = 4).

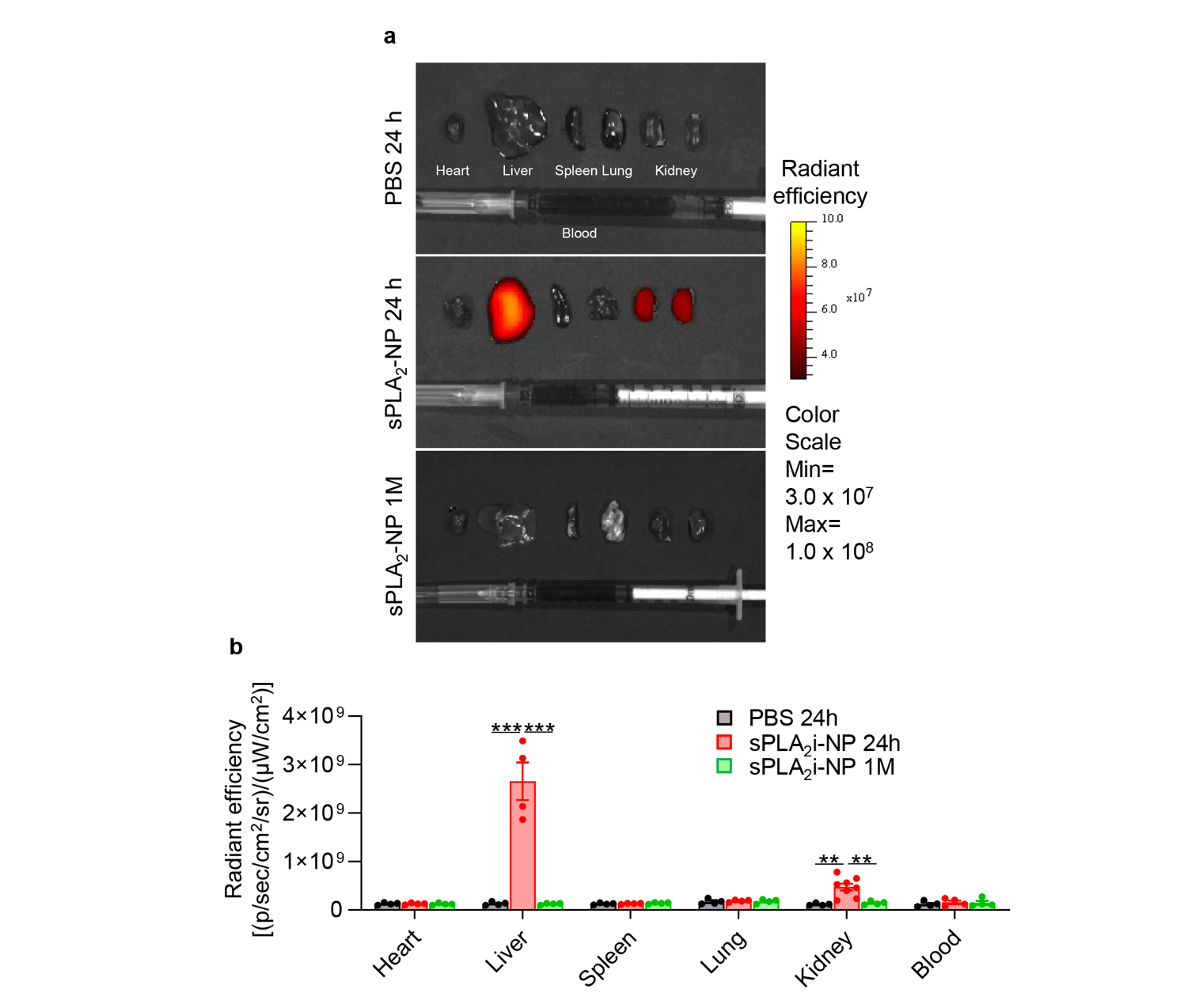

**Supplementary Fig. 7. Systemic biodistribution of sPLA_2_i-NPs following local injection into healthy mouse knee joints. a,** Biodistribution of Cy7-labeled sPLA_2_i-NPs within several internal major organs and blood sample at 24 hours or 1 month post single injection of PBS or Cy7-labeled sPLA_2_i-NPs. Fluorescent scale: max = 3.0 × 10^7^, min = 1.0 × 10^8^. **b,** Quantification of fluorescent radiant efficiency within different organs and blood sample at indicated time points (n = 4). Statistical analysis was performed using one-way ANOVA with Turkey’s post hoc test. Data presented as mean ± s.e.m. **p<0.01, ***p<0.001.

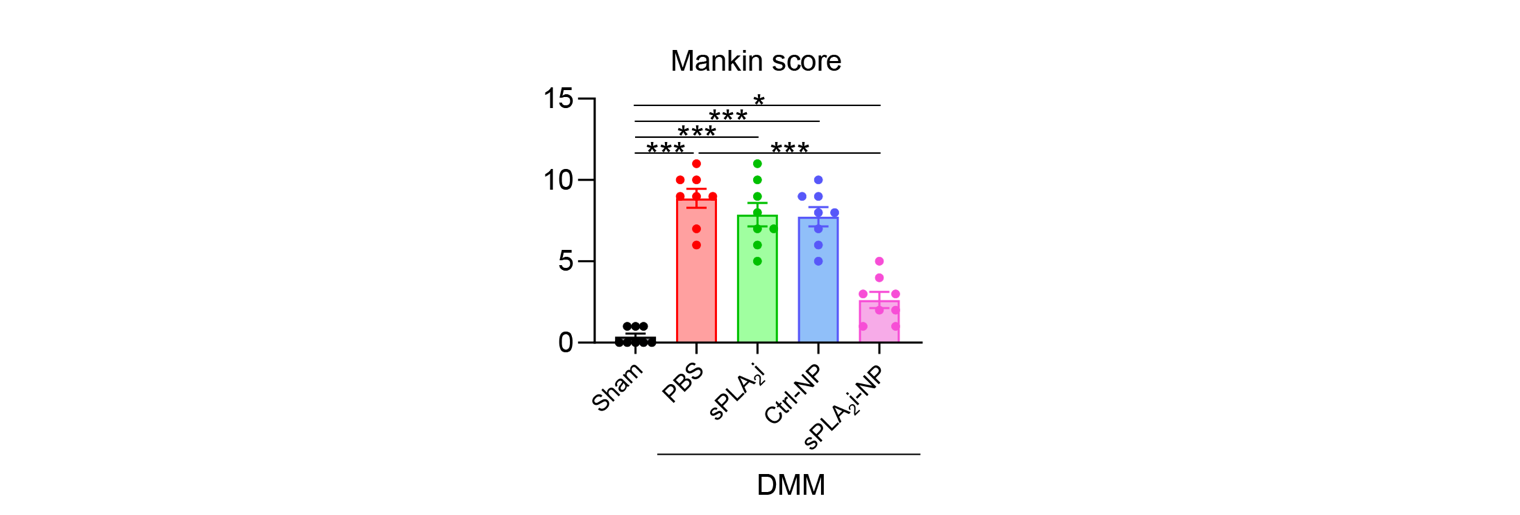

**Supplementary Fig. 8. Effects of sPLA_2_i-NPs on OA progression after 4-month treatment.** The OA severity was accessed by Mankin score after 4-month treatment (n = 8). DMM surgery was performed on 3-month-old male mice followed by intra-articular injections of PBS, sPLA_2_i, Ctrl-NPs and sPLA_2_i-NPs once every week for 4 months. Statistical analysis was performed using one-way ANOVA with Turkey’s post hoc test. Data presented as mean ± S.E.M. *p<0.05, ***p<0.001.

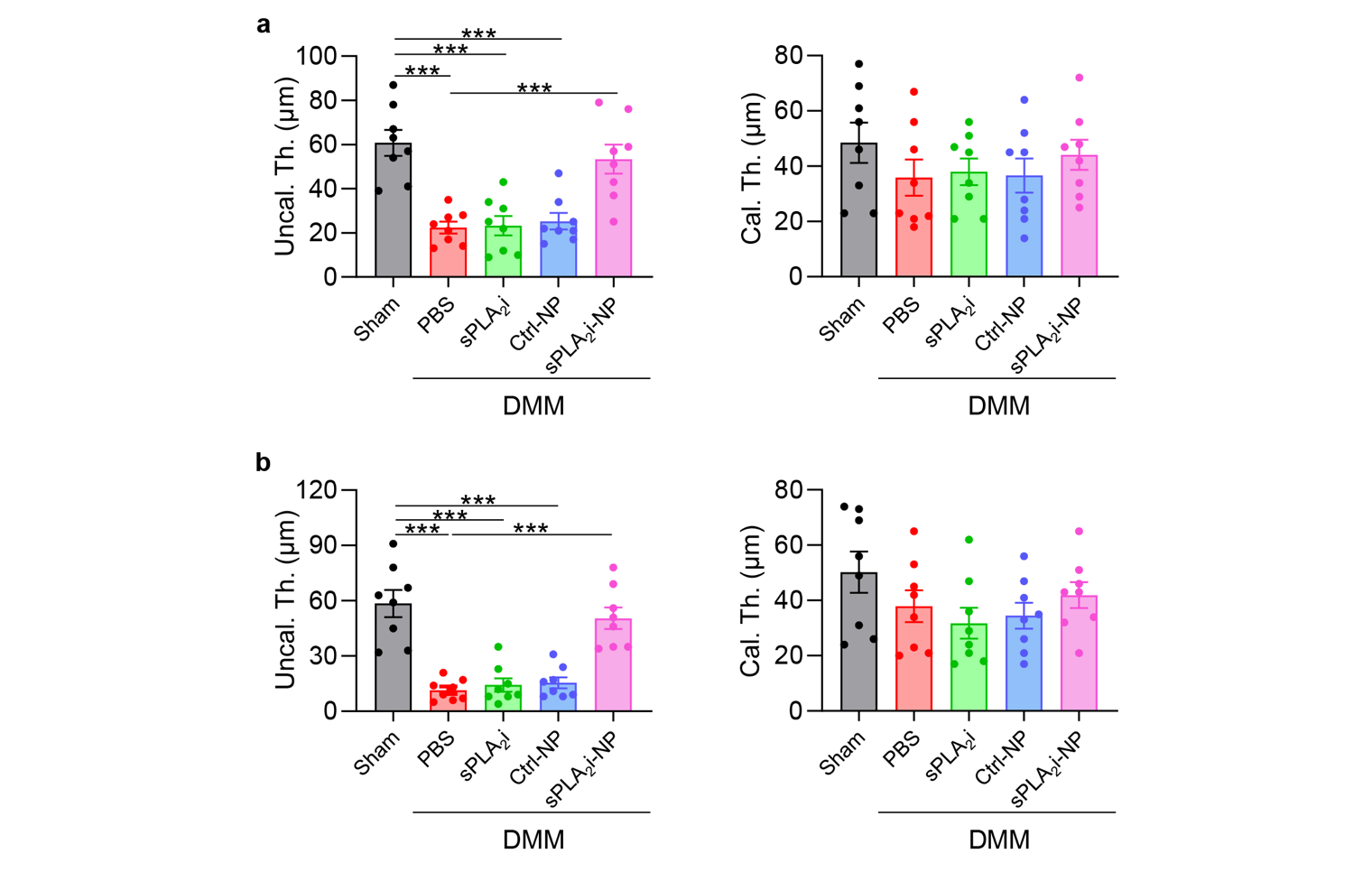

**Supplementary Fig. 9. Effects of sPLA_2_i-NPs on cartilage thickness after 2- or 4-month treatment. a,** Quantification of average thicknesses of uncalcified (Uncal. Th.) and calcified (Cal. Th.) cartilage after 2-month treatment (n = 8). **b,** Quantification of average thicknesses of uncalcified (Uncal. Th.) and calcified (Cal. Th.) cartilage after 4-month treatment (n = 8). DMM surgery was performed on 3-month-old male mice followed by intra-articular injections of PBS, sPLA_2_i, Ctrl-NPs and sPLA_2_i-NPs once every week for 2 or 4 months. Statistical analysis was performed using one-way ANOVA with Turkey’s post hoc test. Data presented as mean ± s.e.m. ***p<0.001.

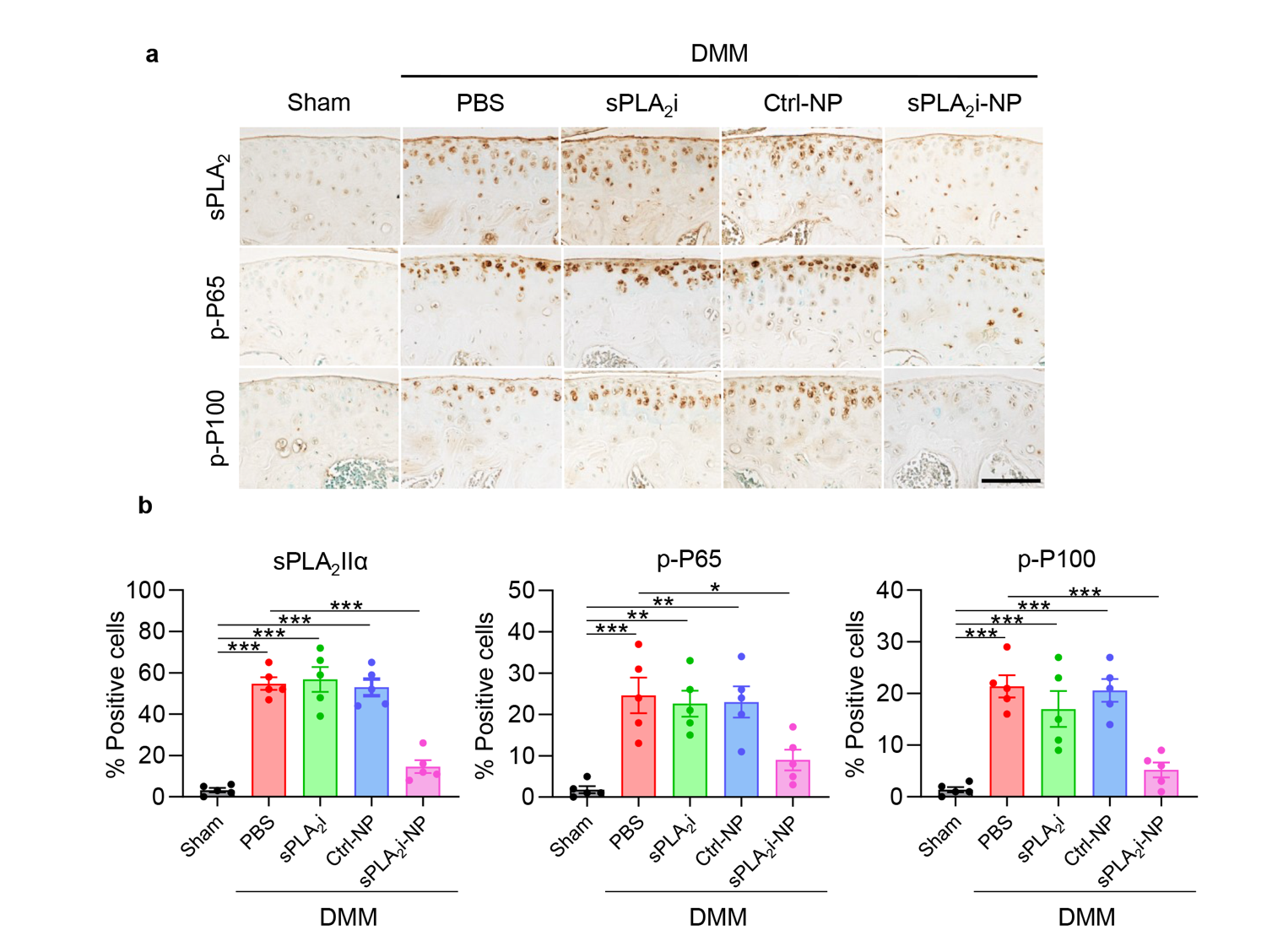

**Supplementary Fig. 10.** **Effects of sPLA_2_i-NPs on altering joint inflammation after 4-month treatment. a,** Representative images of immunohistochemistry staining of sPLA_2_-IIa, p-P65 and p-P100 in the tibial articular cartilage from sham- and DMM-operated knee joints with 4-month treatment. Scale bar, 100 μm. **b,** Quantification of sPLA_2_-IIa-, p-P65- and p-P100-positive chondrocytes in the tibial articular cartilage after 4-month treatment (n = 5). DMM surgery was performed on 3-month-old male mice followed by intra-articular injections of PBS, sPLA_2_i, Ctrl-NPs and sPLA_2_i-NPs once every week for 4 months. Statistical analysis was performed using one-way ANOVA with Turkey’s post hoc test. Data presented as mean ± s.e.m. *p<0.05, **p<0.01, ***p<0.001.

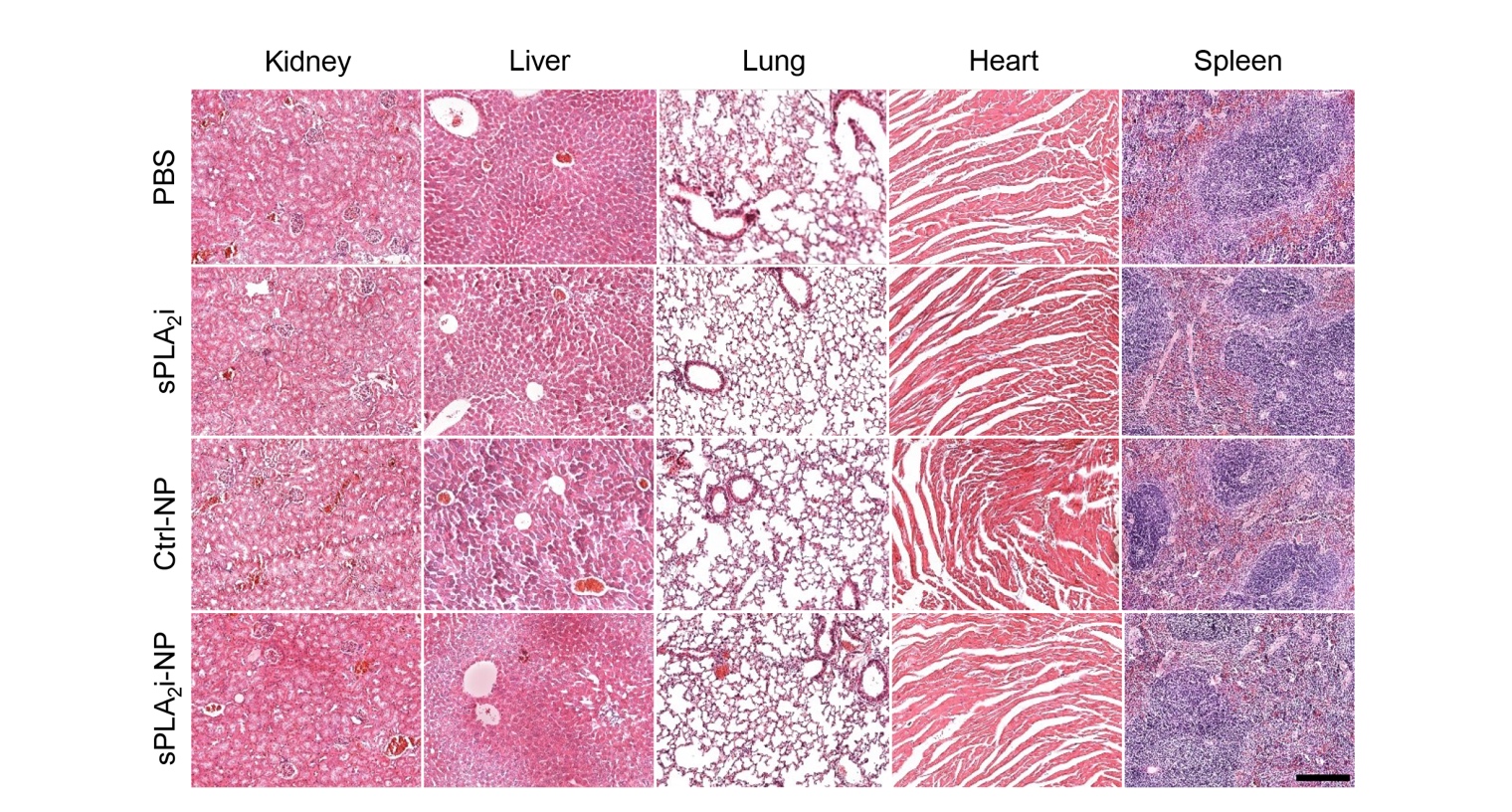

**Supplementary Fig. 11. Histological evaluation of systemic toxicity in vivo.** Representative images of H&E staining of mouse kidney, liver, lung, heart and spleen after 2-month serial injections (once every other week) of PBS, sPLA_2_i, Ctrl-NPs and sPLA_2_i-NPs. Scale bar, 200 μm.

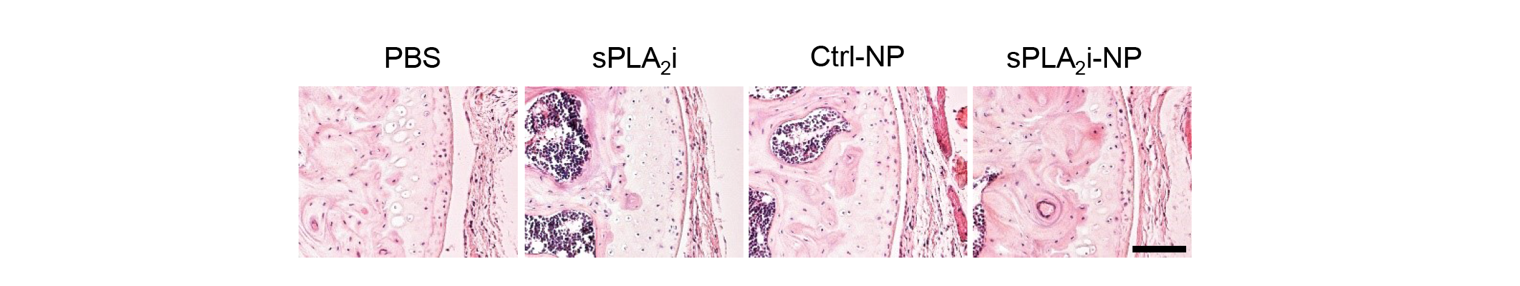

**Supplementary Fig. 12. Histological evaluation of local toxicity in vivo.** Representative images of H&E staining of mouse knee joints after 2-month serial injections (once every other week) of PBS, sPLA_2_i, Ctrl-NPs and sPLA_2_i-NPs. Scale bar, 200 μm.

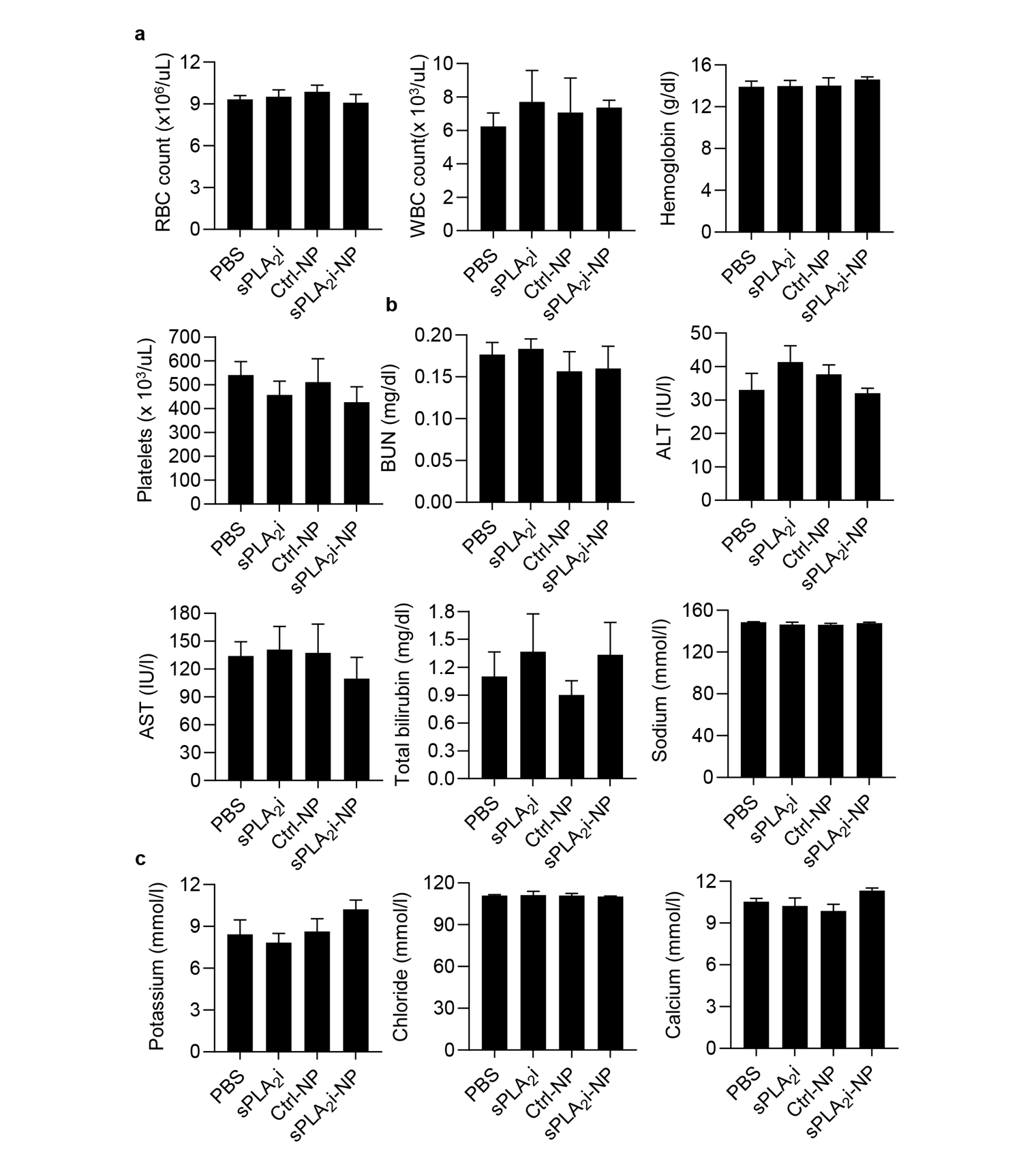

**Supplementary Fig. 13. Serum toxicology makers after 2-month serial injections (once every other week) of PBS, sPLA_2_i, Ctrl-NPs and sPLA_2_i-NPs.** **a,** Hematologic parameters: RBC counts, WBC counts, hemoglobin and platelet counts. **b,** Renal (BUN) and liver (ALT, AST, Total bilirubin) function. **c,** serum electrolytes (sodium, potassium, Chloride and Calcium) showed no significant difference among the 4 treatment groups (n = 3). Statistical analysis was performed using one-way ANOVA with Turkey’s post hoc test. Data presented as mean ± s.e.m.

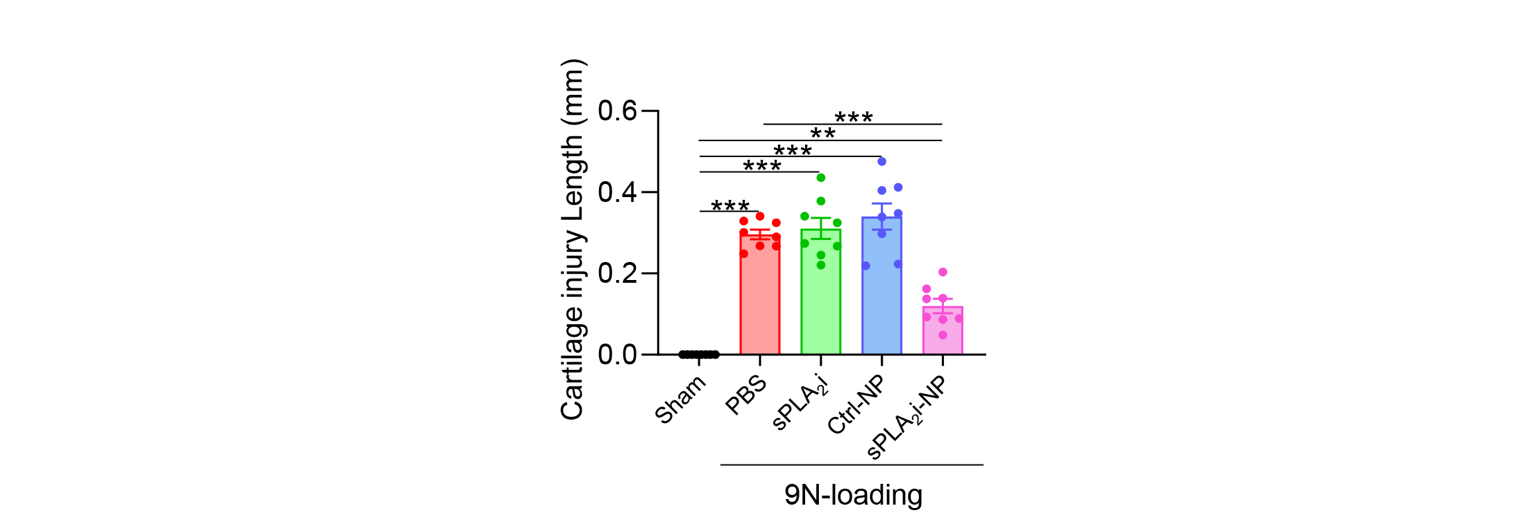

**Supplementary Fig. 14.** **Effects of sPLA_2_i-NPs on acute cartilage injury after 9N-loading.** Quantitative analysis of the length of the cartilage lesion range in the sham- or 9N-load-operated knee joints with 14-day treatment. 9N-loading was performed on 3-month-old male mice. Intra-articular injections of PBS, sPLA_2_i, Ctrl-NPs and sPLA_2_i-NPs were made immediately and 48 hours post loading. Statistical analysis was performed using one-way ANOVA with Turkey’s post hoc test. Data presented as mean ± s.e.m. **p<0.01, ***p<0.001.

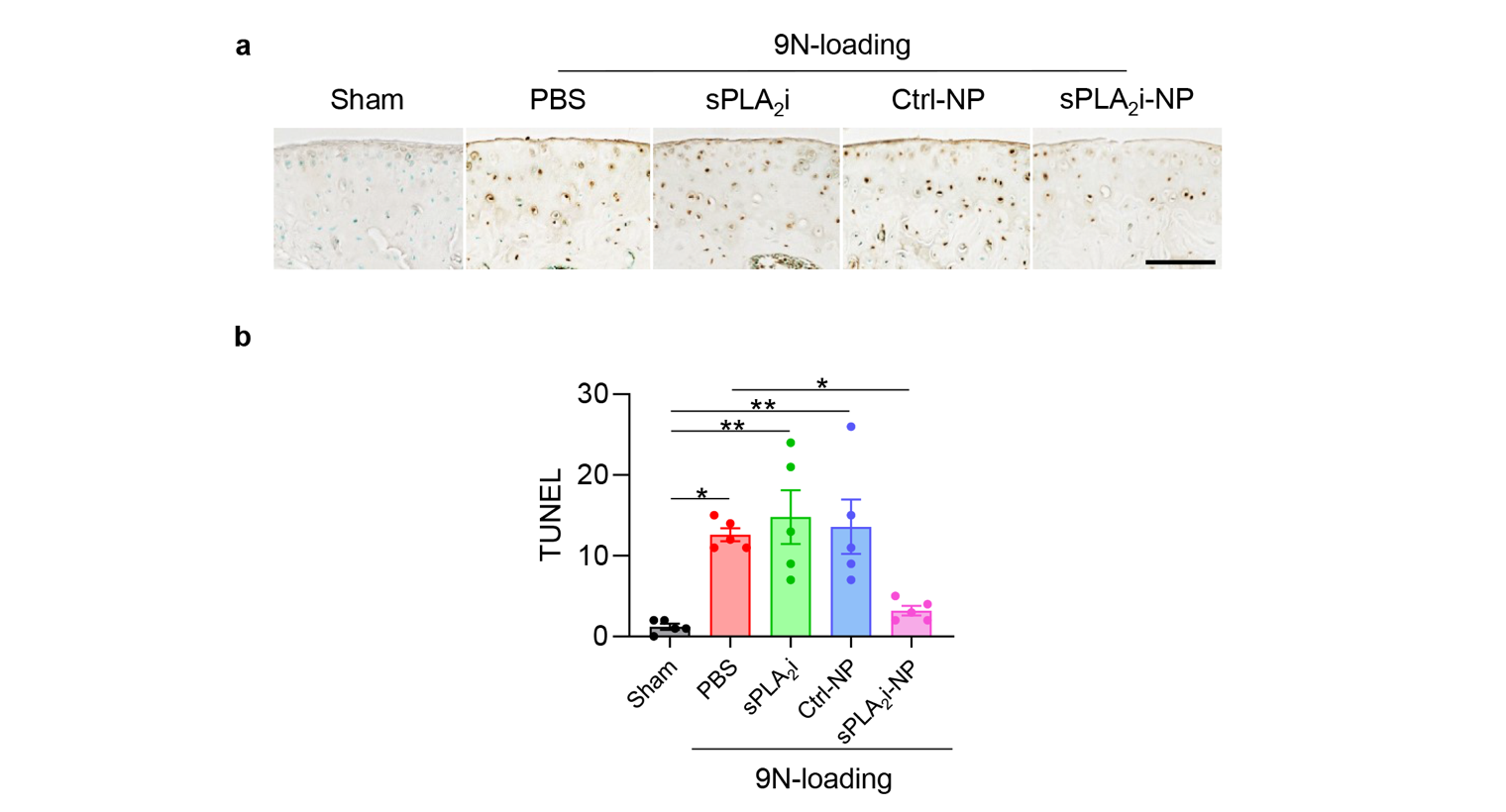

**Supplementary Fig. 15. Effects of sPLA_2_i-NPs on chondrocyte survival after 9N-loading. a,** Representative images of TUNEL staining in the tibial articular cartilage from sham- and 9N-load-operated knee joints with 14-day treatment. Scale bar, 100 μm. **b,** Quantification of TUNEL-positive chondrocytes in the tibial articular cartilage after 14-day treatment (n = 5). 9N-loading was performed on 3-month-old male mice. Intra-articular injections of PBS, sPLA_2_i, Ctrl-NPs and sPLA_2_i-NPs were made immediately and 48 hours post loading. Statistical analysis was performed using one-way ANOVA with Turkey’s post hoc test. Data presented as mean ± s.e.m. *p<0.05, **p<0.01.

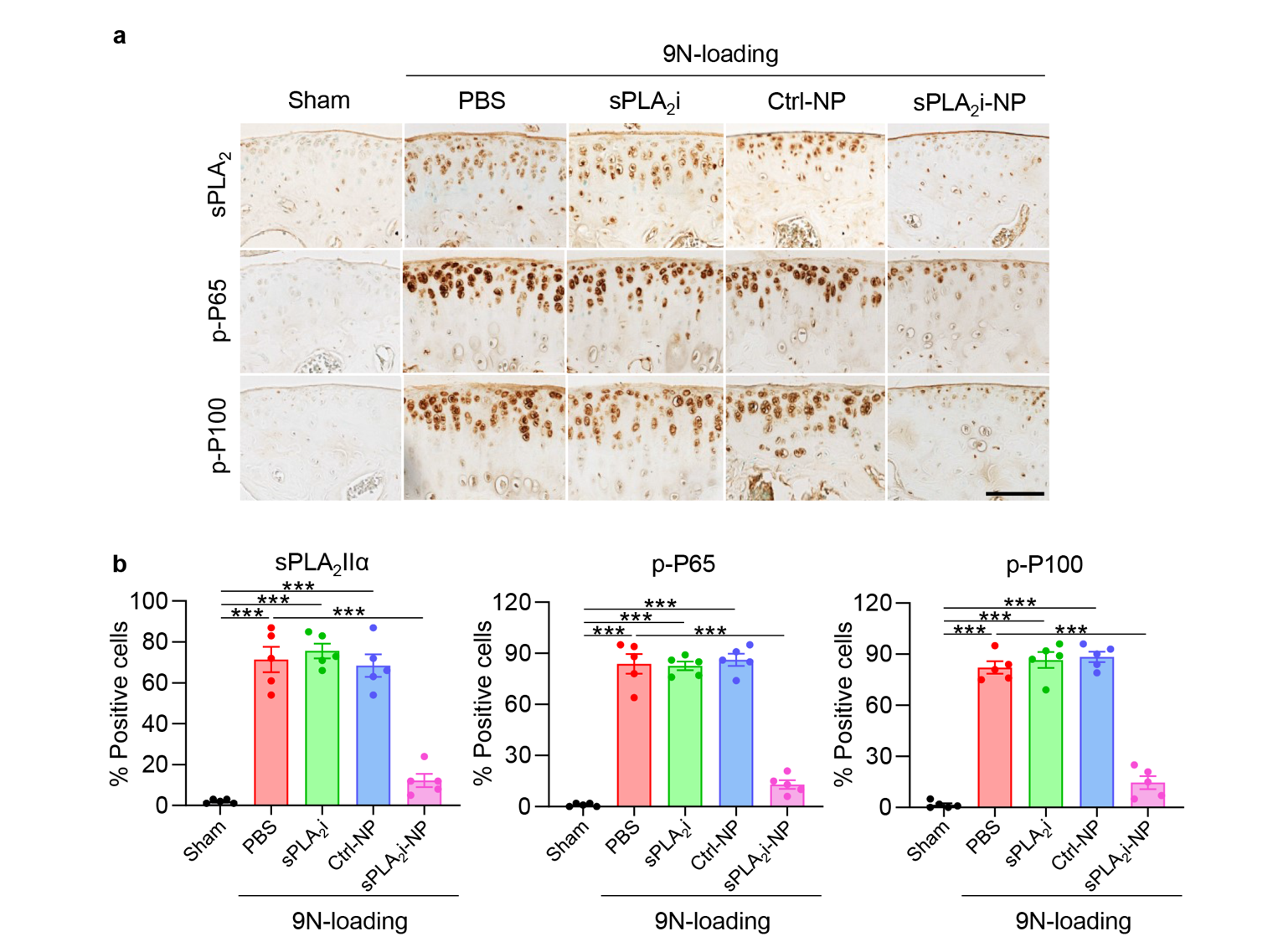

**Supplementary Fig. 16.** **Effects of sPLA_2_i-NPs on altering joint inflammation after 9N-loading. a,** Representative images of immunohistochemistry staining of sPLA_2_-IIa, p-P65 and p-P100 in the tibial articular cartilage from sham- and 9N-load-operated knee joints with 14-day treatment. Scale bar, 100 μm. **b,** Quantification of sPLA_2_-IIa-, p-P65- and p-P100-positive chondrocytes in the tibial articular cartilage after 14-day treatment (n = 5). 9N-loading was performed on 3-month-old male mice. Intra-articular injections of PBS, sPLA_2_i, Ctrl-NPs and sPLA_2_i-NPs were made immediately and 48 hours post loading. Statistical analysis was performed using one-way ANOVA with Turkey’s post hoc test. Data presented as mean ± s.e.m. ***p<0.001.

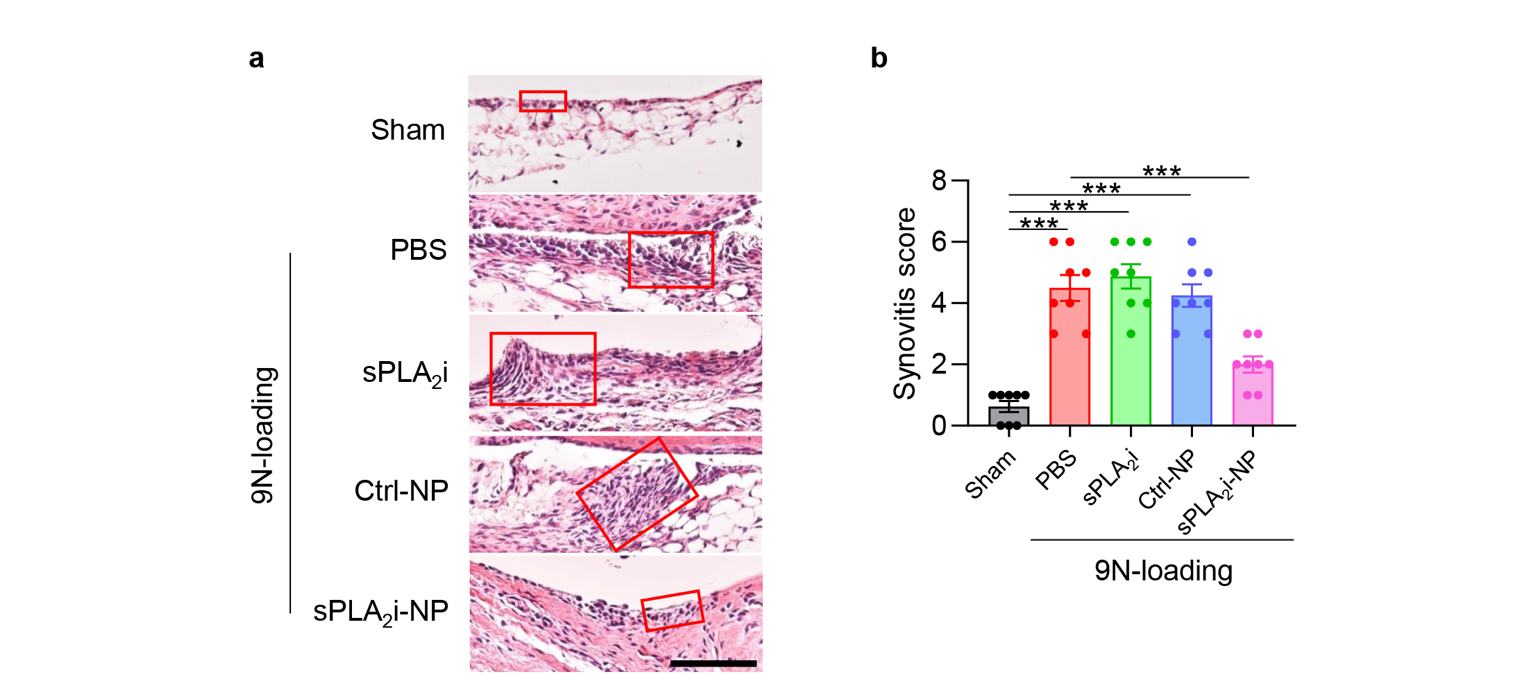

**Supplementary Fig. 17.** **Effects of sPLA_2_i-NPs on altering synovitis after 9N-loading. a,** Representative images of H&E staining in the sham- and 9N-load-operated knee joints with 14-day treatment. Scale bar, 100 μm. Red boxed area indicate the enlargement of synovial lining cell layer. **b,** Synovial inflammation was evaluated by synovitis score after 14-day treatment (n = 8). 9N-loading was performed on 3-month-old male mice. Intra-articular injections of PBS, sPLA_2_i, Ctrl-NPs and sPLA_2_i-NPs were made immediately and 48 hours post loading. Statistical analysis was performed using one-way ANOVA with Turkey’s post hoc test. Data presented as mean ± s.e.m. ***p<0.001.

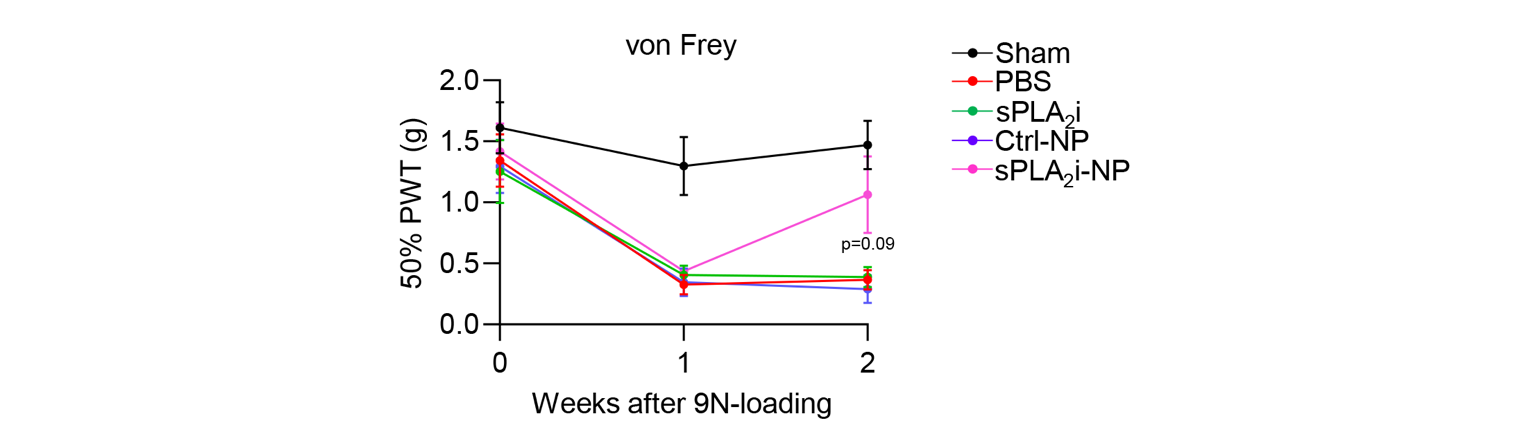

**Supplementary Fig. 18. Effects of sPLA_2_i-NPs on attenuating OA pain after 9N-loading.** von Frey assay on sham and 9N-load-operated knee joints with 14-day treatment at 1 or 2 weeks post loading (n = 8). The data of day 0 was acquired before loading. 9N-loading was performed on 3-month-old male mice. Intra-articular injections of PBS, sPLA_2_i, Ctrl-NPs and sPLA_2_i-NPs were made immediately and 48 hours post loading. Statistical analysis was performed using one-way ANOVA with Turkey’s post hoc test. Data presented as mean ± s.e.m. *p<0.05 for sPLA_2_i-NPs vs. PBS

| Gene | Forward primer | Reverse primer |
| --- | --- | --- |
| sPLA_2_ | 5’-TGGCCTTTGGCTCAATAC-3’ | 5’-GGCATCCATAGAAGGCATAG-3’ |
| Aggrecan | 5’-CTACCGCTGTGAAGTGATG-3’ | 5’-GGTGTAGCGTGTGGAAATAG-3’ |
| Col2a1 | 5’-CAAGAACAGCAACGAGTACCG-3’ | 5’-GTCACTGGTCAACTCCAGCAC-3’ |
| Mmp13 | 5’-TGACCTCCACAGTTGACAGG-3’ | 5’-ATCAGGCACTCCACATCTTGG-3’ |
| Adamts5 | 5’-GCATCCCAGCATTAGGAATTCA-3’ | 5’-GGTGAGAGCTGCATTGGAGGTA-3’ |
| β-actin | 5’-TCCTCCTGAGCGCAAGTACTCT-3’ | 5’-CGGACTCATCGTACTCCTGCTT-3’ |

**Table S1.** Mouse real-time PCR primer sequences used in this study.
